# Allele-specific expression is associated with lower translational efficiency in mouse embryos and stem cells

**DOI:** 10.64898/2026.09.08.750229

**Authors:** Dayea Park, Angélica Liechti, David Gatfield, Can Cenik

**Author notes:** Corresponding author: (CC).

## Abstract

An mRNA’s cytoplasmic fate can be shaped by its transcriptional history, its modifications and associated protein complexes. Allele-specific expression (ASE), in which a gene’s two alleles are transcribed unequally, provides a natural experiment given that the alleles can differ in sequence or chromatin state, yet their transcripts share the same cytoplasm. We previously found lower translational efficiency (TE) among genes with ASE in early mouse embryos. Because that observation came from one cross direction and from preimplantation stages only, sequence-dependent and parent-of-origin-dependent ASE could not be separated, and whether the association held in other cell types remained unknown. Here we addressed both by generating RNA sequencing and ribosome profiling data from reciprocal-cross mouse 8-cell embryos, individual embryonic stem cell clones, neural progenitor cells, and adult tissues. In 8-cell embryos, both sequence-dependent and parent-of-origin-dependent ASE were associated with lower TE. Genes with ASE at the 8-cell stage also had lower protein abundance at the subsequent morula stage. Ribo-ITP, which enables ribosome profiling from single clones, allowed assessing the relationship between ASE and TE in individual embryonic stem cell clones. We found that TE was lower wherever a gene was allelically imbalanced, and the reduction was larger where that imbalance was consistent across clones. The same association held in neural progenitors, and when ASE was defined from independent published datasets, but was not detected in adult tissues. Allelic imbalance in transcription is therefore accompanied by reduced translation of the resulting mRNAs, though the intermediate carrying allelic information to the translation machinery remains unidentified.

## INTRODUCTION

In eukaryotic cells, transcription and translation occur in distinct compartments, yet the two are functionally interconnected. Co-transcriptional RNA processing influences mRNA stability, nuclear export, and translation efficiency (TE) (1–7). Chromatin state and RNA polymerase II elongation kinetics influence these by affecting splice site choice and the recruitment of RNA-binding proteins (5,8,9). For example, the exon junction complex is deposited on mRNAs during splicing and recruiting translation regulatory factors (1,10,11). These findings suggest mRNA transcriptional history shapes its cytoplasmic fate including translation (4,12).

Allele-specific expression (ASE), in which a gene’s two parental alleles are transcribed unequally, provides a natural test of this idea. Alleles may differ stably, through cis-regulatory variation or allele-specific chromatin state (13–16), or expression may fluctuate stochastically without any stable difference between them (17–19). Consequently, transcripts originating from the two alleles may differ in processing dynamics, RNA-binding protein occupancy, and modification state.

However, whether these allele-specific nuclear histories affect translation has been little studied, despite extensive characterization of ASE at the transcriptional level, particularly during early embryogenesis. The most extreme form of ASE is imprinting, in which one allele is silenced in a parent-of-origin-dependent manner (20). Beyond imprinting, widespread allelic biases are observed throughout preimplantation development and across embryonic lineages (15,21,22). Given that gene regulatory programs are still being established at this stage, these biases are both prevalent and variable. Yet allele-specific differences in RNA abundance need not be reflected proportionally at the protein level, because protein output also depends on translational regulation (23–25).

We previously observed that genes with ASE had lower translation efficiency than genes without ASE in 4-and 8-cell embryos obtained from a single cross direction of C57BL/6J (B6) × CAST/EiJ (CAST) mouse (26). Given the lack of reciprocal crosses or experiments in other cell types, several key questions remained open including: (1) Is reduced TE associated with both sequence-dependent and parent-of-origin-dependent ASE? (2) Is the association general or restricted to particular cell types? (3) Does the relationship depend on the consistency of allelic bias across cells? In the current study, we address these questions by generating RNA-seq and ribosome profiling data from 8-cell embryos of reciprocal B6 × CAST crosses, which separate sequence-dependent from parent-of-origin-dependent ASE, and from mouse embryonic stem cells (mESCs), mESC-derived neural progenitor cells (NPCs), and adult kidney, liver, and lung.

Given that ASE varies between individual cells (27–29), addressing these questions requires clonal measurements of translation to separate stable from stochastic allelic bias. Such measurements have until recently been infeasible owing to input requirements, a challenge that we overcome here by using Ribo-ITP (ribosome profiling via isotachophoresis), which enables analysis from single embryos and single stem cell clones. We find that both classes of ASE are accompanied by reduced TE in 8-cell embryos, that the association persists in mESCs and NPCs but is not detected in adult tissues. Finally, we find that within clones, the TE reduction is strongest for genes with consistent allelic bias.

## RESULTS

### Identifying ASE in preimplantation mouse embryos using reciprocal crosses

In our previous study using hybrid mice derived from B6 × CAST crosses, we quantified allele-specific read counts in mouse embryos from zygote to the 8-cell embryos using RNA-seq reads that uniquely map to SNPs distinguishing the two genomes (26). Here, we used the same set of 210,005 SNPs between these genomes to enable transcriptome-wide quantification of allelic RNA-seq counts from a newly generated RNA-seq data obtained from 8-cell stage embryos of the reciprocal cross (CAST × B6) (Fig 1A, S1 Fig). Allelic read counts were reproducible between replicates at both the individual SNP and gene levels (Spearman’s correlation > 0.6; S2 Fig). For example, *Reep1* contains 20 SNPs, all of which were detected as showing B6 allele biased expression. Similarly, *Smim48* contains 12 SNPs, and all exhibited a consistent bias toward the CAST allele (Fig 1B).

**Fig 1.**
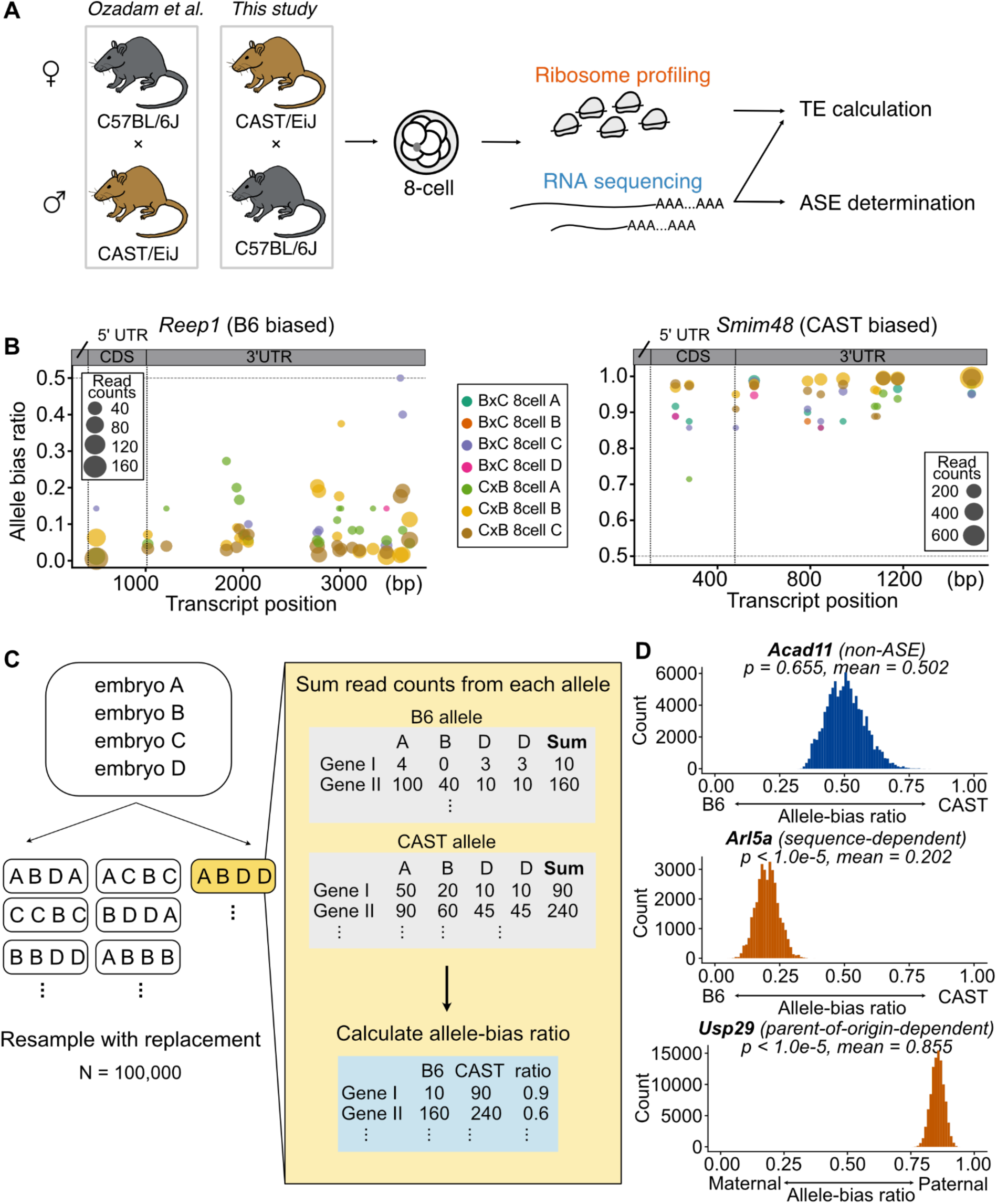
Defining ASE from mouse reciprocal cross. (A) Schematic overview of the experimental and analytical workflow. RNA-seq and ribosome profiling were performed on individual 8-cell embryos from reciprocal B6×CAST (BxC) and CAST×B6 (CxB) crosses to quantify ASE and TE. (B) Allelic read counts and allele-bias ratios at individual SNP sites. The y-axis shows the allele-bias ratio, calculated as the number of CAST allele reads divided by the total number of B6 and CAST allele reads at each SNP site. The x-axis indicates the genomic position of each SNP within the gene. Dot size represents the read count at each SNP, and colors denote individual reciprocal 8-cell embryos. The left panel shows *Reep1*, which exhibits B6 allele-biased expression, whereas the right panel shows *Smim48*, which exhibits CAST allele-biased expression. (C) Schematic illustration of the bootstrap-based method used to define ASE. For each bootstrap iteration, reciprocal embryo samples were resampled with replacement while preserving the original sample size (e.g., if 7 reciprocal embryo samples were available, 7 samples were resampled with replacement). The four embryos shown are a simplified illustration of the resampling procedure. (D) Representative bootstrap output. The top panel shows a gene without ASE, where the distribution of resampled allele-bias ratios is centered near 0.5. The middle panel shows a gene with sequence-dependent ASE, with preferential expression of the B6 allele. The bottom panel shows a gene with parent-of-origin-dependent ASE, with preferential expression of the paternal allele. For sequence-dependent analyses, allele-bias ratios were calculated as CAST/(B6 + CAST). For parent-of-origin analyses, allele-bias ratios were calculated as paternal/(maternal + paternal), after assigning allelic reads according to parental origin in each reciprocal cross. P-value and mean ratio is calculated from bootstrap resampling.

Because allele-specific read quantification depends on the generation and analysis of RNA-seq data, we next assessed the influence of experimental and computational approaches on these measurements. We first compared two independent RNA-seq library preparation methods (Smart-seq3 and NEB; Methods) and found that gene-level allelic read counts were consistent between the two methods (Spearman’s correlation > 0.72; S3 Fig). We then examined the effect of the computational analysis pipeline by comparing GATK with our in-house custom pipeline (26,30). Similarly, allelic read counts were consistent between the two computational pipelines.

Across both the Smart-seq3 and NEB datasets, gene-level allelic read counts quantified using GATK and our in-house SNP counting pipeline showed a Spearman’s correlation > 0.92 (S4 Fig). Together, these results indicate that although experimental and computational choices introduce some variability in allele-specific read quantification, the overall measurements remain comparable.

The allele-specific RNA-seq read counts were used to identify genes exhibiting ASE in 8-cell stage embryos using a bootstrap-based approach (Fig 1C, Methods). Because RNA-seq data were generated from reciprocal crosses, we could distinguish two classes of ASE: sequence-dependent ASE, in which allelic bias follows strain identity regardless of parental origin, and parent-of-origin-dependent ASE, in which allelic bias follows parental origin across reciprocal crosses. Overall, we identified 86 sequence-dependent and 168 parent-of-origin-dependent genes with ASE. For example, *Acad11* was classified as non-ASE (mean allele-bias ratio = 0.502; p = 0.655). In contrast, *Arl5a* exhibited a strong and consistent B6 allele bias and was classified as a gene with sequence-dependent ASE (mean allele-bias ratio = 0.202; p < 1.0e-5). Similarly, *Usp29* exhibited a strong and consistent paternal bias and was classified as a gene with parent-of-origin-dependent ASE (mean paternal allele-bias ratio = 0.855; p < 1.0e-5, Fig 1D). Among imprinted genes defined from two previous reports (31,32), we detected two genes as sequence-dependent ASE (*Gab1* and *Impact*) and two as parent-of-origin-dependent ASE (*Snx14* and *Usp29*) (Fig 1C). Although *Gab1* and *Impact* have been reported as imprinted, evidence supporting their imprinting during early embryonic development remains limited. In contrast, *Usp29* is an established paternally expressed imprinted gene in mice (33). Its paternal expression in our 8-cell embryos was consistent with previous reports (Fig 1D). No gene ontology terms were significantly enriched among genes with ASE, as their annotated functions span diverse cellular functions, including the cell cycle, metabolism, signal transduction, transport, and transcriptional regulation. Together, these analyses established a high-confidence set of genes showing sequence-and parent-of-origin-dependent ASE for subsequent analyses.

### Both sequence-and parent-of-origin-dependent ASE in 8-cell mouse embryos associate with reduced TE

The method of TE calculation from ribosome profiling and RNA-seq data has been debated (34). To ensure that our results are robust to the method of choice, throughout the study we used two methods: a compositional regression model (compositional TE) (35) and the ribosome to RNA ratio (ratio TE) (Methods). Because ratio TE is known to be confounded by differences in mRNA abundance (36), we compared the RNA expression of the genes defined as ASE at the 8-cell stage across embryonic development stages. We found that they had significantly higher expression levels, with median RPKM differences ranging from 0.04 to 13.14 across 2-to 16-cell stages (Fig 2A). This observation may appear counterintuitive, as silencing of one allele might be expected to reduce overall RNA abundance. One possible explanation is that genes with low expression generate fewer allele-informative reads, reducing the statistical power to detect ASE and potentially biasing detection toward more highly expressed genes. To minimize potential confounding arising from RNA expression differences, we ensured that all comparisons in the rest of the manuscript used a randomly selected subset of genes without ASE such that the distribution of their RNA expression is matched with that of genes with ASE unless otherwise noted (Methods).

**Fig 2.**
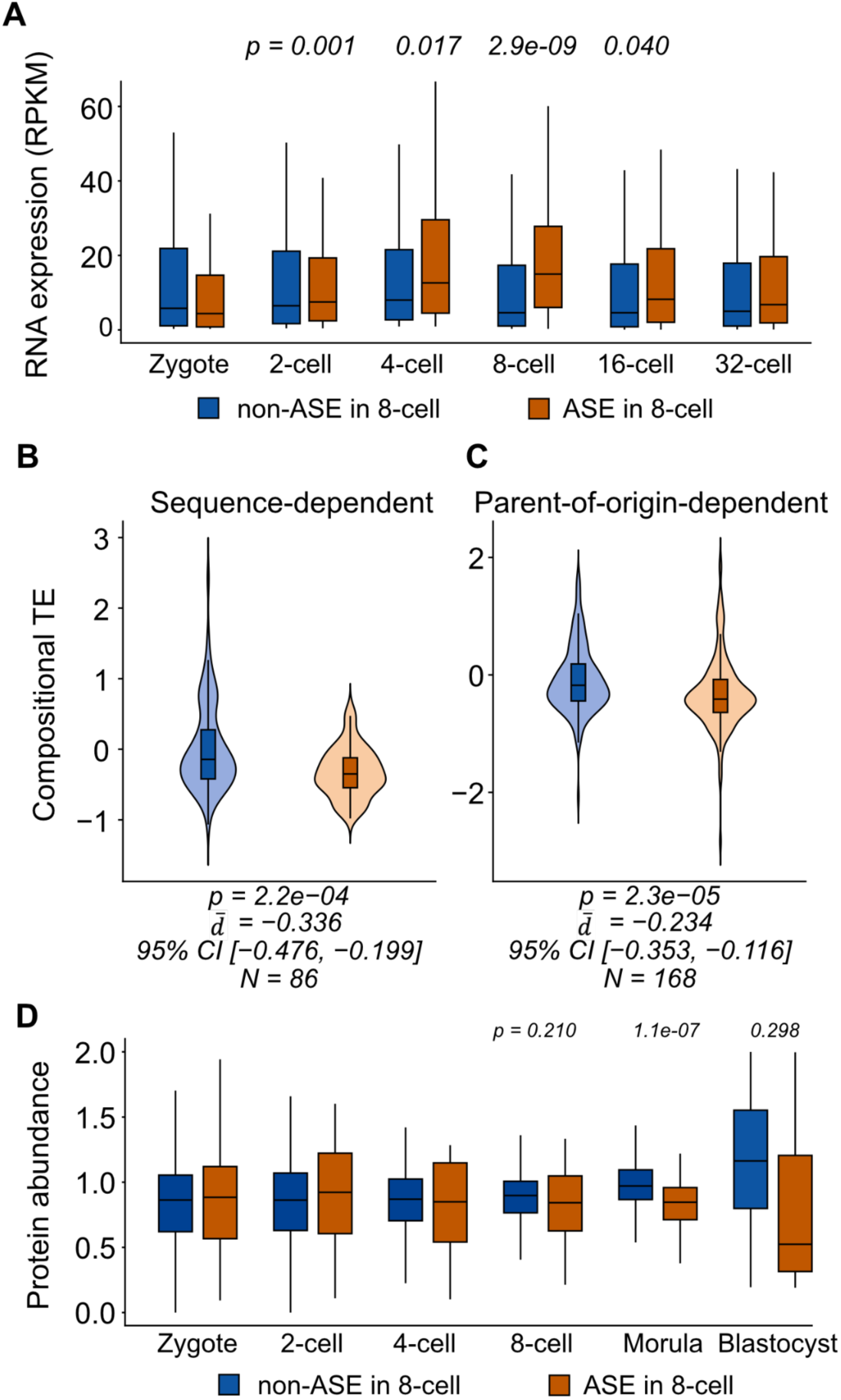
Genes with ASE exhibit reduced translation efficiency in 8-cell stage mouse embryos. (A) Comparison of mRNA abundance between genes defined as ASE and non-ASE across mouse preimplantation development. mRNA abundance at the zygote to 8-cell stages was obtained from Ozadam et al. (2023) and quantified as Reads Per Kilobase of transcript per Million mapped reads (RPKM). mRNA abundance at the 16-and 32-cell stages was obtained from Ghatpande et al. (2025). (B–C) Comparison of compositional TE between genes with and without ASE (B, sequence-dependent ASE; C, parent-of-origin-dependent ASE). Statistical significance was assessed using two-sided Wilcoxon rank-sum tests (*p*). The 95% CIs for the mean TE difference were estimated by nonparametric bootstrap resampling within each group using the 2.5th and 97.5th percentiles. Effect sizes represent the mean TE difference between genes with and without ASE (*TE_ASE_* – *TE_NON-ASE_*). *N* indicates the number of genes with ASE included in each comparison. (D) Comparison of protein abundance between genes defined as ASE and non-ASE across mouse preimplantation development. Protein abundance data were obtained from Gao et al. (2017).

In our previous study, we observed that genes displaying ASE were translated significantly less efficiently at both the 4-cell and 8-cell stages (26). Given that major zygotic genome activation has been completed and the maternally inherited transcripts are largely depleted by the 8-cell stage (37,38), we focused on this stage to assess whether our initial observation is robustly detected in reciprocal crosses. We generated ribosome profiling in addition to RNA-seq data allowing us to assess whether the reduced TE of genes with ASE is reproducible.

Importantly, reciprocal crosses enabled us to distinguish sequence-and parent-of-origin-dependent ASE, allowing us to determine whether these distinct forms of allelic regulation differ in their association with TE. We found that genes exhibiting either sequence-or parent-of-origin-dependent ASE exhibited reduced TE. The mean compositional TE differences were −0.336 for sequence-dependent ASE and −0.234 for parent-of-origin-dependent ASE (p =2.2e−04 and 2.3e−05, respectively), indicating that both classes of genes exhibiting ASE exhibit lower TE than genes without ASE. The corresponding 95% confidence intervals (CI) were entirely below zero (−0.476 to −0.199 and −0.353 to −0.116, respectively; Fig 2B,C). Similar results were observed when using ratio TE (S5 Fig). Consistent with the technical robustness observed at the allelic read counts and total RNA-seq counts (Spearman correlation > 0.87, S3 Fig), reduced TE in genes with ASE is reproducibly detected across different library preparations (Smart-seq3 vs NEB; S6 Fig).

We next wondered if the reduced TE among genes with ASE is reflected in their corresponding protein abundance (24,25,39). Hence, we analyzed mass spectrometry data from mouse preimplantation development from the zygote to blastocyst stages (40). In particular, the ribosome occupancy in 4-cell and 8-cell embryos had been shown to be highly correlated with protein abundance in the subsequent morula-stage of development (26). Despite having on average higher mRNA expression than genes without ASE, we found that the corresponding protein abundance of genes with ASE was significantly reduced in the morula stage (Fig 2D). The discrepancy between RNA and protein levels is consistent with post-transcriptional differences limiting protein production from transcripts with ASE.

Taken together, reciprocal cross analysis enabled identification of ASE and its inheritance patterns. We found that both sequence-and parent-of-origin-dependent ASE are associated with reduced TE. Notably, although 8-cell stage genes with ASE exhibit elevated RNA expression on average during early embryogenesis, they show reduced protein abundance at later stages. These findings suggest that translational repression of genes with ASE may limit protein production in early mouse development.

### Cell type-specific association between ASE and reduced TE

The reduction in TE associated with ASE in 8-cell stage embryos raises the question of whether this relationship is specific to early development or is also observed in other cell types and tissues. To address this, we generated matched RNA-seq and ribosome profiling data from individual clones of mouse embryonic stem cells (mESCs), mESC derived neural progenitor cells (NPCs), and adult kidney, liver, and lung tissues all from B6 × CAST hybrid mice (Fig 3A).

**Fig 3.**
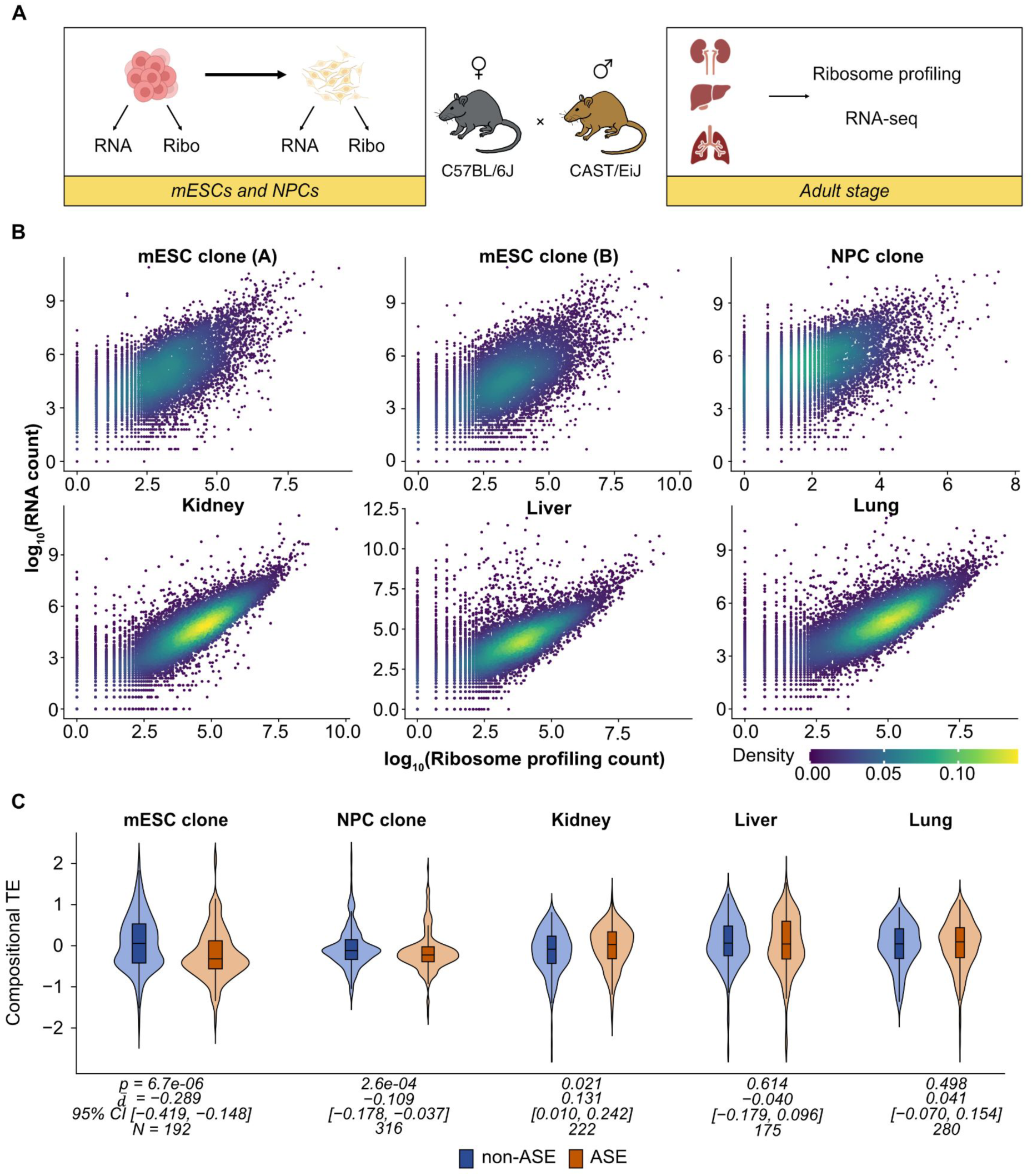
ASE is associated with lower TE in mESCs and NPCs. (A) Experimental design to investigate TE across developmental stages and tissues. Individual clones of mESCs and NPCs were collected, and matched RNA-seq and ribosome profiling libraries were generated from the same clone. For the adult stage, kidney, liver, and lung tissues were collected, and matched RNA-seq and ribosome profiling data were generated from biological replicate bulk tissue samples. (B) Correlation between RNA-seq and ribosome profiling gene counts across two individual mESC clones, one NPC clone, and one replicate each from kidney, liver, and lung samples. Spearman’s correlation coefficients were calculated using log10-transformed RNA-seq (y-axis) and ribosome profiling (x-axis) read counts. Points represent individual genes and are colored according to local point density, estimated using two-dimensional kernel density estimation. (C) Comparison of compositional TE between genes with and without ASE in mESC individual clones, kidney, liver, and lung. Statistical significance was assessed using two-sided Wilcoxon rank-sum tests (*p*). The 95% CIs for the mean TE difference were estimated by nonparametric bootstrap resampling within each group using the 2.5th and 97.5th percentiles. Effect sizes represent the mean TE difference between genes with and without ASE (*TE_ASE_*-*TE_NON-ASE_)*. *N* indicates the number of genes exhibiting ASE included in each comparison.

Before comparing ASE and TE, we first evaluated the quality and reproducibility of the mESC and NPC data. All RNA-seq and ribosome profiling libraries had strong replicate-to-replicate correlations (Spearman’s correlation > 0.9; Fig 3; S7 Fig). In addition, all ribosome profiling libraries had expected read-length distributions, metagene profiles, and enrichment of reads mapping to coding regions (S8 Fig).

Having established the technical robustness of these datasets, we then determined ASE from RNA-seq and estimated TE using the same approach used for 8-cell embryos. Across mESC clones, we identified 192 genes with ASE and these had lower TE (compositional TE difference−0.289, p = 6.7e−06; Fig 3C). For NPCs, we first confirmed successful differentiation by qRT-PCR analysis of marker genes and by comparison to previously published transcriptomic data (S9 Fig) (41). Among NPCs, we identified 314 genes with ASE. TE was also reduced among genes with ASE, but the effect size was smaller (compositional TE difference −0.109, p = 2.6e−04; Fig 3C). Taken together these results indicate the association between ASE and reduced TE is similarly observed among the individual clones of both mESCs and NPCs.

Given the smaller effect size observed in NPCs than in mESCs, we next examined differentiated adult tissues to determine whether the association between ASE and reduced TE is maintained or lost. We observed high technical quality of experiments (S10 Fig), and identified 222, 175, and 280 genes showing ASE in kidney, liver and lung, respectively. In contrast to 8-cell embryos and mESCs, genes with ASE did not exhibit reduced compositional TE in these tissues (kidney, TE difference 0.131, p = 0.021; liver, TE difference −0.040, p = 0.614; lung, TE difference 0.041, p = 0.498; Fig 3C). The same results were observed when using ratio TE (S11 Fig). Together, these results indicate that the association between ASE and reduced TE is strongest in preimplantation embryos, and mESCs, detectable in NPCs with a smaller effect size, and is largely absent in differentiated adult tissues.

### Consistent ASE across clones is associated with stronger translational repression

Allelic ratios for a given gene are heterogeneous across cells and embryos (41,42). To assess the heterogeneity of allele-specific expression across individual samples, we calculated the mean allelic ratio and its standard deviation across 8-cell embryos and mESC clones for each gene (Fig 4A). Given this heterogeneity, on average, 151 out of the 192 genes classified as ASE exhibited allelic bias in a given mESC clone (S12 Table).

**Fig 4.**
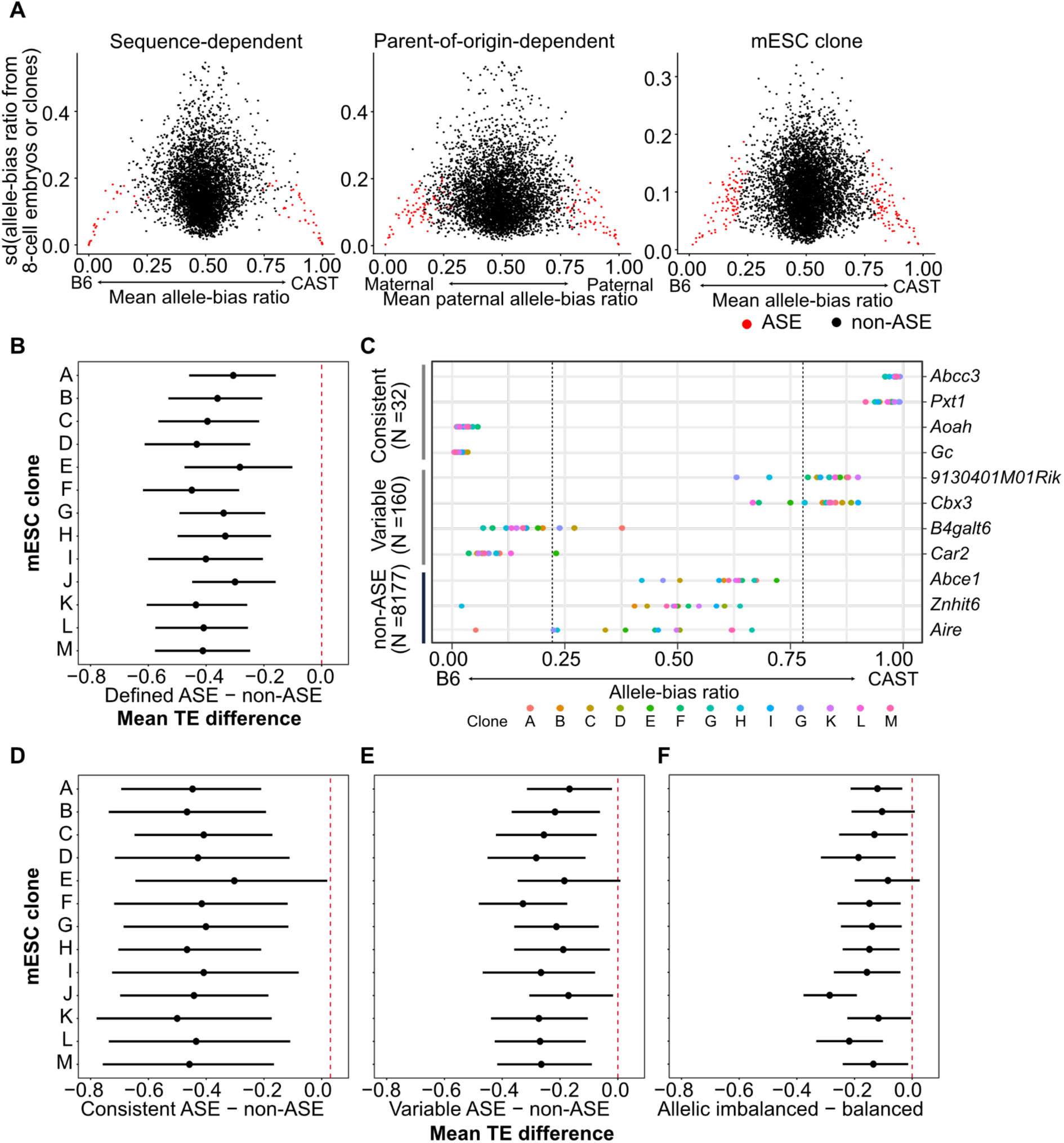
Genes with consistent ASE exhibit greater reduction in TE. (A) Heterogeneity of allele-bias ratios across individual 8-cell embryos and mESC clones. The mean (x-axis) and the standard deviation (y-axis) of allele-bias ratios across embryos or clones, representing the degree of ASE heterogeneity were plotted. For reciprocal-cross 8-cell embryo datasets, heterogeneity was calculated separately for sequence-dependent (left) and parent-of-origin (middle) allele-bias ratios. Heterogeneity across mESC clones is shown in the right panel. Red dots indicate genes defined as ASE in our analysis. (B,D-F) Clone-specific differences in compositional TE between genes with and without allelic bias, calculated as *TE*_’*with allice*_ *_bias_* – *TE* _’*without allice*_ *_bias_* Comparisons were performed separately within each mESC clone. Dots indicate the mean TE difference, and error bars represent 95% confidence intervals. Comparisons are shown between genes with ASE and genes without ASE (B), genes with consistent ASE and genes without ASE (D), genes with variable ASE and genes without ASE (E), and genes classified as non-ASE overall but exhibiting clone-specific allelic imbalance versus genes without allelic imbalance in the corresponding clone (F). (C) Classification of consistent, variable ASE and non-ASE. Consistent ASE genes showed ASE (allele-bias ratio < 0.2 or > 0.8) in the same direction across all informative clones, with measurements available in more than five clones. Genes not meeting these criteria were classified as variable ASE. Colors represent individual mESC clones, and plots show allele-bias ratios across clones.

To test whether this heterogeneity in ASE across clones relates to the observed difference in translation, we next compared TE between groups of genes within each clone while matching for RNA expression. As expected, the subset of genes with ASE that exhibit significant allelic bias in a given clone exhibited lower TE compared to genes without ASE in that clone (mean of compositional TE differences from −0.44 to −0.28 across clones; p < 0.001, Fig 4B; S13 Table; ratio TE S13A Fig).

We then classified genes with ASE into two groups, consistent and variable ASE. Genes were considered to have consistent ASE when the allele-bias ratio met the predefined threshold (ratio < 0.2 or > 0.8) across all informative clones, whereas genes were considered to have variable ASE when the threshold was met in some clones but not others (Fig 4C; S14 Fig; Methods). In mESCs, 32 genes met the definition of consistent ASE while the remaining 160 were deemed to have variable ASE.

Genes with consistent ASE exhibited a substantially greater reduction in compositional TE relative to genes without ASE (mean of compositional TE difference from −0.50 to −0.30 across clones; p < 0.05 in all clones; Fig 4D; S13 Table). Genes with variable ASE also showed reduced TE, but with a smaller effect size (mean compositional TE difference from −0.32 to −0.16 across clones; p < 0.05 in all clones; Fig 4E; S13 Table). Ratio TE estimates showed the same overall pattern, with a stronger reduction among genes exhibiting consistent ASE (S15 Fig).

Given that genes with consistent ASE also had larger magnitude allelic imbalance, we next asked whether progressively stronger allelic bias is associated with progressively lower TE. We grouped each gene with ASE in each clone using progressively more stringent allelic ratio thresholds. Compositional TE was subsequently compared across these groups. Genes meeting more stringent thresholds exhibited progressively lower TE (S16 Fig).

Heterogeneity in allelic bias across clones also allows a test of whether reduced TE reflects properties of the genes classified as ASE or the allelic state of a gene in a particular clone. If TE tracks allelic state, genes not classified as ASE across the dataset should nonetheless show reduced TE in clones where they are allelically imbalanced. Using clone-level allelic ratios, we identified genes with allelic imbalance in individual clones that did not meet the bootstrap significance threshold for ASE hence classified as non-ASE when considering all clones. Compared with RNA expression-matched genes with no allelic imbalance in the same clone, these genes showed reduced TE in every clone (mean compositional TE difference −0.28 to −0.08; p < 0.10 in all clones; Fig 4F, S13 Table). Together, these results show that reduced TE accompanies allelic imbalance within individual clones and that the reduction is strongest where allelic bias is consistent across clones.

### Consistent ASE across independent datasets is associated with reduced TE

Our clonal analysis used a single mESC line, and a modest sample size of 13 clones. The set of genes that are detected as ASE varies between laboratories and mESC batches. Agreement between independent studies therefore provides a second axis of consistency, complementary to consistency across clones within our dataset. To exploit this, we used published datasets that report allelic expression despite lacking matched ribosome profiling. Hence, ASE calls were derived from the published data, while TE was measured using our own libraries.

We first reanalyzed a reciprocal 8-cell embryo dataset generated from the same cross (43). Using our bootstrap approach, we identified 112 genes with ASE across 11 embryos, 31 of which were also classified as ASE in our dataset. The lowest TE was observed for genes classified as ASE in both datasets (compositional TE difference = −0.500, *p* = 0.021; Fig 5A; ratio TE in S17 Fig), suggesting that consistent ASE across independent datasets is associated with stronger translational reduction. To reduce potential confounding from RNA expression levels and expression-dependent ASE detection power, the ASE and non-ASE groups were matched for RNA expression in each comparison.

**Fig 5.**
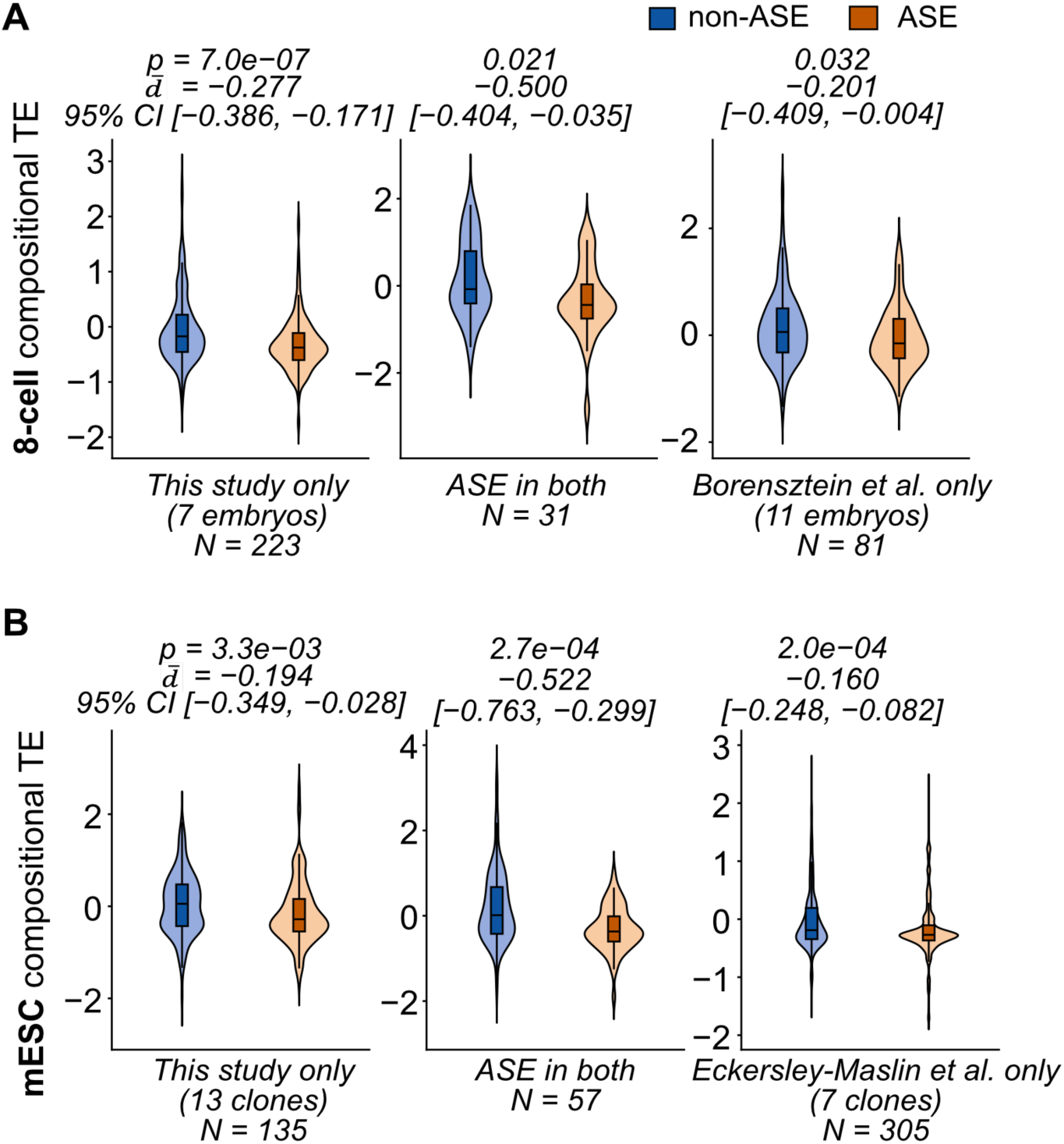
TE comparison between genes with ASE across studies. (A-B) Comparison of TE between genes with and without ASE using independent 8-cell embryo (A) and mESC (B) datasets. The left panels compare the compositional TE of genes exhibiting ASE identified in this study. The right panel shows the TE comparison in independent datasets (8-cell embryos: Borensztein et al.; mESCs: Eckersley-Maslin et al.). The middle panel compares TE between genes without ASE and genes consistently identified as ASE in both datasets.

Using a similar approach, we examined published mESC (41). After reanalysis, we identified 362 with ASE in the published dataset and 57 genes exhibited ASE in both datasets. Compared with genes identified as ASE in only one dataset, these shared genes showed the greatest reduction in compositional TE (mean compositional TE difference −0.522; Fig 5B). A similar pattern was observed using ratio TE (S18 Fig). Together, these observations suggest that greatest reduction in TE was observed for genes identified as exhibiting ASE across studies, further supporting the relationship between consistent ASE and lower TE.

## DISCUSSION

Allele-specific expression is accompanied by lower translational efficiency, and this holds for both sequence-dependent and parent-of-origin-dependent ASE (Fig 2C,D). It remains unclear whether this finding reflects a shared molecular pathway or convergent outcomes of distinct processes. Several features of our analysis argue that this association is not an artifact of a single analytical or experimental choice. Reduced TE among genes with ASE was recovered using two independent quantifications of translational efficiency (a compositional regression model and a ribosome-to-RNA ratio), and in independently generated datasets. The direction of the effect was consistent across embryos, mESCs, and NPCs, was retained after matching genes with and without ASE on mRNA abundance, and was insensitive both to library preparation chemistry and to the SNP-calling pipeline used to assign allelic reads.

In this study, we found that genes with ASE showed lower protein levels at the morula stage (Fig 2D). This finding is consistent with the observation that ribosome occupancy at the 4-and 8-cell stages has been shown to be highly correlated with protein abundance at the subsequent morula stage (26). Genes with ASE have higher mRNA levels on average (Fig 2A), so transcript abundance alone would predict higher rather than lower protein expression. Hence, the reduction in protein abundance therefore occurs in spite of a transcriptional difference in the opposite direction. These findings suggest that lower translation efficiency of genes with ASE may limit protein production in early embryogenesis.

Resolving the relationship between ASE and TE within clones required matched RNA-seq and ribosome profiling from individual mESC clones, a measurement that has not previously been feasible due to input limitations for ribosome occupancy measurements. We overcame this challenge with Ribo-ITP. Despite substantial heterogeneity in allelic bias (27), reduced TE was preserved within individual clones (Fig 4B). Consistent ASE, which typically reflects stable genetic or epigenetic differences between alleles, showed a stronger reduction in TE, whereas variable ASE, which likely reflects stochastic fluctuations, showed a weaker effect. Translational differences therefore track stable allelic states.

The relationship also varied with cellular context. It was strongest in preimplantation embryos and mESCs, contexts marked by high dosage sensitivity and extensive epigenetic reprogramming (44,45), weaker in NPCs, and undetectable in adult kidney, liver, and lung. We are cautious about interpreting this gradient. The tissue comparison differs from the others in several ways. The samples were bulk rather than clonal, were prepared using a different ribosome profiling protocol, and required a different operational definition of ASE. Any of these could reduce sensitivity independently of biology, and cell-type-specific relationships could be masked by averaging across the heterogeneous cell populations that constitute a tissue. What our data suggest is that the association is not a universal property of ASE and determining where it does apply will require allele-resolved transcriptomics combined with low-input translational profiling in defined differentiated cell types.

What could link allelic differences in expression to reduced translation? The most direct possibility is that sequence differences between the alleles alter translation themselves. For example, cis-variants can change RNA secondary structure, uORF presence, codon composition, and thereby affect ribosome recruitment and elongation. Such a mechanism cannot provide a general explanation, however, given that parent-of-origin-dependent ASE, where the allelic bias reverses between reciprocal crosses, was also strongly associated with reduced TE.

An alternative mechanism may involve differential modification of the mRNAs transcribed from the two alleles. m6A has been proposed to modulate ribosome recruitment and translation efficiency (46,47), raising the possibility that allele-specific m6A could differentiate transcripts originating from the two alleles. Yet we observed no enrichment or depletion of allele-specific m6A among genes with ASE (S19 Fig) (48), and only two genes previously identified as carrying allele-specific m6A exhibited ASE. The small overlap between confidently called allele-specific m6A events and genes with ASE precludes us from unambiguously distinguishing an absence of effect from a lack of statistical power. However, our observations argue that allele-specific m6A is unlikely to be sufficient to explain the correlation between ASE and TE.

Consequently, we have not yet identified a mechanism to account for the observed relationship. The framing that motivated this work posits that allele-specific nuclear histories are transmitted to the cytoplasm through RNA-binding protein occupancy, or modification state. In this work, we have support at the level of the outcome for this framework but not at the level of mechanism. The exon junction complex (EJC) offers a precedent for this kind of gap. Spliced mRNAs were known to be translated more efficiently well before that difference could be mechanistically explained (1,49). While EJC was initially characterized as a splicing-dependent mark with roles in nuclear export and surveillance (50,51), its recruitment of factors acting directly on the translation machinery was established later (10). The relationship we describe may prove similar, awaiting identification of the intermediate that carries allelic information from the nucleus to the translation machinery.

The central limitation of this study is that every result is correlative. We have not shown that allelic imbalance causes reduced translation, and we have not identified a mechanism that would. A direct test requires inducing or removing ASE at a defined locus and measuring the translational consequence, which in turn requires allele-specific perturbation. We attempted this by designing single guide RNAs against promoter sequences containing SNPs, with the aim of modulating one allele selectively, but were unable to establish sufficient allelic specificity to draw any conclusions. Until allele-specific perturbation can be coupled to an allele-resolved translational readout, the direction of causality and whether the two are causally connected at all will remain untested.

All of the comparisons above are between genes. Separating allele-specific translational regulation from measurement noise will require approaches that decouple allele assignment from footprint length. One promising direction would be to leverage Ribo-STAMP or related methods combined with long-read sequencing (52,53) ideally paired with allele-resolved proteomics to establish whether allelic differences in ribosome occupancy produce allelic differences in protein. Taken together, our results support a robust association between allelic differences in transcript abundance and lower TE during early mouse development, most strongly where ASE is stable across clones. The relationship is reproducible across methodologies, independent datasets, and cell contexts, but its cause and mechanism are unresolved.

## MATERIAL AND METHOD

### Mouse embryo collection

Experiments involving preimplantation mouse embryos were approved by the Institutional Animal Care and Use Committee at the University of Texas at Austin (AUP-2022-00114). We obtained C57BL/6J × CAST/EiJ (B6 × CAST) F1 embryo data from a previous study (26). To analyze embryos from the reciprocal cross, we collected CAST × B6 F1 8-cell embryos using the same procedures described in Ozadam et al. 2023. Individual CAST × B6 8-cell embryos were collected in ITP lysis buffer (20 mM BisTris titrated to pH 7.2 with HCl, 1.0% Triton-X100, 5mM MgCl2, 2.5 mM CaCl_2_, 100 mM NaCl, 1 mM DTT, 0.1 mg/ml cycloheximide), and used for ribosome profiling and RNA-seq library preparations.

### Cell culture

The B6 × CAST F1 hybrid mESCs were kindly provided by Dr. David Spector (54). Cell culture and neurodifferentiation of mESCs were conducted following previously described protocols (41,48). To derive NPCs, mESCs were cultured in a 1:1 mixture of DMEM/F12 (Sigma) and Neurobasal medium (Gibco), supplemented with 1× N2 (Gibco), 1× B27 (Gibco), 40 mg/L insulin (Gibco), and 25 μg/mL BSA fraction V (Sigma). Cells were seeded at 1 × 10⁵ cells per well in a 6-well plate and maintained for 6 days with daily medium changes. Subsequently, cells were resuspended in N2 expansion medium (DMEM/F12 containing 1× N2, 50 μg/mL BSA fraction V [Sigma], 10 ng/mL epidermal growth factor [PeproTech], 10 ng/mL fibroblast growth factor [PeproTech], and 1 μg/mL laminin [SouthernBiotech]) and plated onto uncoated T75 flasks to form neurospheres. After 4 days, neurospheres were collected by centrifugation at 200 × g for 3 minutes and replated on gelatin-coated plates in N2 expansion medium. After two to three passages, the cultures yielded a homogeneous population of NPCs. Single mESC and NPC clones were then collected by colony picking for library preparation.

### Quantitative real-time PCR

To validate successful differentiation, the expression of mESC (*Nanog*, *Pou5f1*, and *Sox2*) and NPC marker genes (*Tubb3*, *Map2* and *Nes*) was assessed by quantitative real-time PCR (qRT-PCR). Total RNA was isolated from NPCs using Direct-zol RNA kit (Zymo Research) and reverse transcribed into cDNA using SuperScript IV Reverse Transcriptase (Thermo Fisher Scientific) according to the manufacturer’s instructions. qRT-PCR was performed using PowerUp SYBR Green Master Mix (Thermo Fisher Scientific) on a QuantStudio 7 Flex Real-Time PCR System (Thermo Fisher Scientific). Each 20 μL reaction contained 10 μL of 2× PowerUp SYBR Green Master Mix, 100 ng of cDNA, 1 μM each forward and reverse primer, and nuclease-free water to a final volume of 20 μL. Amplification was performed using the manufacturer’s recommended cycling conditions: 50°C for 2 min, 95°C for 2 min, followed by 40 cycles of 95°C for 15 s and 60°C for 1 min. A melt curve analysis was performed following amplification to verify the specificity of the PCR products. Relative gene expression was normalized to *Actb* and calculated using the 2^−ΔΔCt^ method. Primer sequences are provided in the S20 Table. Statistical analyses were performed using data from three biological replicates, each with three technical replicates.

### Single clone selection for library preparation

mESCs and NPCs were seeded at low density (<2,000 cells) in a 10 cm dish. When a single cell formed a colony, the cells were washed once with 1× PBS containing 0.1 mg/ml cycloheximide. A single clone was then picked under a microscope (10×) in less than 2 µL and collected in 6 µL of pre-lysis buffer (0.1% Triton X-100, 1 mg/mL cycloheximide in water). After pipetting the sample up and down 10 times, half of the lysate was used for RNA-seq and the remaining half for ribosome profiling library preparation by adding 1 µL of 5× lysis buffer (100 mM Bis-Tris, pH 7.2, 25 mM MgCl₂, 25 mM CaCl₂, 500 mM NaCl, 5 mM DTT).

### Sample collection from mouse adult tissues

Liver, kidney and lung samples were collected from male F1 hybrid mice originating from B6 × CAST crosses, age 7 weeks. Organ samples were a kind gift from Eric Aeby and Jeannie Lee. Animals were subjected to brief anesthesia (isofluran) before sacrifice by decapitation and organs were immediately extracted, flash frozen in liquid nitrogen, and stored at −80 °C until further processing.

### Ribosome profiling library preparation from 8-cell embryo, mESC and NPC clones

To obtain ribosome footprints from low-input samples such as single embryos and single clones, we performed ribosome profiling via isotachophoresis (Ribo-ITP) as described previously (55). For RNA digestion, 1:50-diluted micrococcal nuclease (MNase; NEB) was used for embryos, and 1:300-diluted RNase I (Ambion) was used for mESC and NPC clones, followed by incubation at 37 °C for 30 minutes for digestion. This process was stopped by adding 1 µL of 10 mM EGTA (for MNase) or 1% SDS (for RNase I), and the lysate was subsequently processed by Ribo-ITP. Ribosome footprints were eluted by collecting RNA fragments between the 19 nt and 36 nt markers.

Ribosome profiling libraries were generated using the D-Plex Small RNA-seq Kit (Diagenode) with minor modifications. The dephosphorylation reaction was supplemented with 0.5 µL T4 PNK (NEB) and incubated for 25 minutes. cDNA was amplified for 8 cycles (clones) or 12 cycles (embryos). The amplified products were cleaned with 1.8× AMPure XP beads, and a ∼200 bp band was excised from a 10% TBE gel. DNA was extracted from the gel by overnight incubation in 300 µL of PAGE-DNA extraction buffer (10 mM magnesium acetate, 0.5 M ammonium acetate, and 1 mM EDTA) on a rotor at room temperature. DNA was precipitated with 600 µL of 100% ethanol and 1.5 µL of GlycoBlue (ThermoFisher) for 1 h at −20 °C, followed by centrifugation at 13,000 × g for 1 h at 4 °C. The DNA pellet was resuspended in nuclease-free water. Pooled libraries were sequenced on an Illumina NovaSeq 6000 and Novaseq X plus platform.

### RNA-seq library preparation from 8-cell embryo, mESC and NPC clones

To prepare RNA-seq library from an 8-cell embryo, two methods were used: Smart-seq3 (version 3) (56) and NEBNext Single Cell/Low Input RNA Library Prep Kit (NEB). Single CAST × B6 8-cell embryos were lysed and reverse transcribed following the modified Smart-seq3 method as described previously (26). cDNA was pre-amplified using 13 PCR cycles and purified with 1.8× AMPure XP beads, followed by elution in 8 µl of nuclease-free water. One microliter of pre-amplified cDNA was quantified using the Qubit double-stranded DNA high-sensitivity assay. Samples were diluted with nuclease-free water, normalized to 1.2 ng input in 6.5 µl, and subjected to tagmentation and post-tagmentation PCR. Tagmentation was performed with 7.5 µl of tagmentation mix, followed by the addition of 9 µl Nextera Index primers and 6× tagmentation PCR mix. Sixteen PCR cycles were conducted, and the products were purified using a 1× AMPure XP ratio. The final library size distribution and concentration were assessed using a High Sensitivity DNA Bioanalyzer, and sequencing was performed on an Illumina NovaSeq 6000.

For libraries prepared using the NEBNext Kit, we followed the manufacturer’s instructions. Cells were lysed in 1× lysis buffer and subjected to three freeze–thaw cycles at −80 °C. Following reverse transcription, cDNA was amplified using 11 PCR cycles for single clones and 18 cycles for single embryos. Amplified cDNA was purified using 0.6× AMPure XP beads. For fragmentation, cDNA input was normalized to 3 ng, and adapter ligation and indexing were performed using the NEBNext Multiplex Dual Oligos kit (NEB) with 8 PCR cycles. Final cleanup was carried out with 0.9× AMPure XP beads. Library quality was assessed using a Bioanalyzer, confirming a narrow fragment size distribution with a peak at 300–350 bp. The sequencing was performed on NovaSeq X Plus.

### Ribosome profiling library preparation from adult tissues

Snap-frozen tissue pieces (∼450 mg per liver, 350 mg per kidney, 200 mg per lung sample) were recovered from −80 °C storage and homogenised in 3 volumes of lysis buffer (150 mM NaCl, 20 mM Tris-HCl pH 7.4, 5 mM MgCl_2_, 5 mM DTT, 1% Triton X-100, 0.5% Sodium deoxycholate, 100 µg·mL^−1^ cycloheximide, complete EDTA-free protease inhibitors (Roche), 40 U·mL^−1^ RNasin plus (Promega)), cleared from debris, quantified via OD_260_, and digested with RNase I (Ambion) and Turbo DNase (Ambion) as described in Arpat et al. (2020). After separation through Sephacryl S-400 HR spin columns (GE Healthcare Life Sciences), Qiazol/miRNeasy RNA extraction kit was used on the column flow-through to obtain purified RNA that contains the ribosome-protected fragments as well as contaminating RNA species. After rRNA depletion via Ribo-Zero Gold rRNA Removal Kit (MRZG12324 Illumina) from 5 µg (liver, lung) or 3 µg (kidney) input RNA, rRNA-depleted RNAs were purified and concentrated (Zymo) and synthetic 30 nt and 60 nt RNA oligos were added to all samples as spike-ins, as previously described (57). Subsequently, ribosome profiling libraries were generated from this material using the TruSeq Ribo-Profile Kit [RPHMR12126 Illumina; all described in Arpat et al. (2020)]. Libraries were sequenced in-house on Illumina HiSeq platform.

### RNA-seq library preparation from adult tissues

Parallel RNA-seq libraries from all tissue samples were prepared essentially following the Illumina protocol; briefly, after total RNA extraction (miRNeasy RNA Extraction Kit, Qiagen) from an aliquot of the same lysates from which the footprints had been generated, ribosomal RNA was depleted with Ribo-Zero Gold rRNA (Illumina), samples were heat-fragmented, the same RNA spike oligos as for ribosome-protected fragment libraries were added, and sequencing libraries generated as reported (58). Libraries were sequenced in-house on Illumina HiSeq platform.

### RNA-seq read processing

RNA-seq read processing was performed using a bash script available on GitHub (rnaseq_process_featurecount.sh). Adaptor trimming was performed using Cutadapt v1.18 (59) with following parameters: “-m 20 --trimmed-only -g ATTGCGCAATG” (Smart-seq3) and “-a AGATCGGAAGAGCACACGTCTGAACTCCAGTCA -A AGATCGGAAGAGCGTCGTGTAGGGAAAGAGTGT -O 8 -m 20 --cores=8” (NEB). Trimmed reads were filtered to remove rRNA and tRNA contaminants using Bowtie2 v2.3.4.3, and the remaining reads were aligned with STAR (v.2.7.10b) using the parameters: “--outSAMmapqUnique 10 --outSAMtype BAM SortedByCoordinate --quantMode GeneCounts --readFilesCommand zcat” (60).

The mouse transcriptome was derived from the GRCm38 genome assembly, with gene annotations from GENCODE vM25 (61). Using our custom Python script (generate_appris_reference.py) developed in a previous study (62), we selected the longest APPRIS principal transcript. For this alignment, an N-masked transcriptome was used as the reference. To minimize alignment bias toward the B6 reference genome, 210,005 SNP positions distinguishing B6 and CAST were masked by replacing the reference bases with ‘N’. The final masked reference transcriptome was generated using a previously described custom script (translate_genomic_to_transcriptomic_coord.ipynb) (26) (S21 Table).

After alignment, duplicate reads were removed using Samtools (samtools rmdup) (63). Gene-level read counts were quantified using featureCounts with the APPRIS mouse GTF annotation (GRCm38 assembly, GENCODEM25), using paired-end (-p) and unstranded (-s 0) counting (64). Raw counts were normalized using a centered log-ratio transformation. Data quality was assessed by calculating pairwise gene-level correlations on the normalized counts using Spearman’s rank correlation, implemented in the stats package (v 4.1.1) in R.

### Ribosome profiling read processing

Ribosome profiling data were processed using RiboFlow v0.0.1 (62,65). Unique Molecular Identifier (UMI) sequences were extracted using UMI-tools with a regex pattern specifying the first 12 nucleotides as the UMI and the following 4 nucleotides for removal (--extract-method=regex). Subsequently, adapter sequences were trimmed using Cutadapt (-a AAAAAAAAAACAAAAAAAAAA --overlap=4 --trimmed-only), requiring a minimum 4-nt overlap and retaining only reads containing the adapter (66). Trimmed reads were first aligned to mouse rRNA and tRNA references using Bowtie2 v2.3.4.3, and unaligned reads were subsequently mapped to the N-masked transcriptome (67). Reads with a mapping quality score greater than two were retained and deduplicated using UMI-tools with the directional adjacency method (--read-length option) (S21 Table). The quality control was performed and ribosome profiling read counts were then extracted from .ribo files using RiboR and Ribograph (62,65) (S8,22 Fig).

### Detection of ASE

Using BAM files generated from RNA-seq alignments to the N-masked transcriptome, allelic read counts were quantified with GATK ASEReadCounter version 4.0 (30). For allele-specific analyses, we considered 210,005 SNPs located within the transcript regions.

Allelic count tables were processed in R. Genes with low coverage were filtered out if they had fewer than 10 reads in more than half of the samples. For example, in a dataset of 8 embryo samples, genes were retained only if they had more than 10 reads in at least five samples. The SNP level allelic counts were summed per gene. Genes located on the X chromosome were excluded from all analyses.

For embryo analyses, allelic counts from reciprocal crosses were analyzed separately to distinguish sequence-dependent from parent-of-origin-dependent ASE. Sequence-dependent ASE was defined by grouping reads by strain origin B6 or CAST across reciprocal crosses. Parent-of-origin-dependent ASE was defined by grouping reads by maternal or paternal origin.

To assess ASE, we applied a bootstrapping strategy adapted from Ozadam et al., 2023 (26). Bootstrapping was used to estimate the robustness of allelic bias across biological replicates while accounting for sampling variability arising from limited replicate numbers and sequencing depth. For each condition, replicate samples were randomly resampled with replacement while preserving the original number of replicates: 8 reciprocal embryos, 13 mESC clones and 5 NPC clones. Reads were then aggregated across replicates. This procedure was repeated 100,000 times. In each iteration, allele-bias ratio was calculated as:

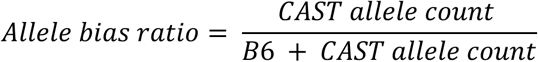

For each gene, the mean allele-bias ratio and empirical p-values were recorded. P-values were calculated separately for each direction of allelic bias and the minimum value was retained. Specifically, one-sided p-values were defined as:

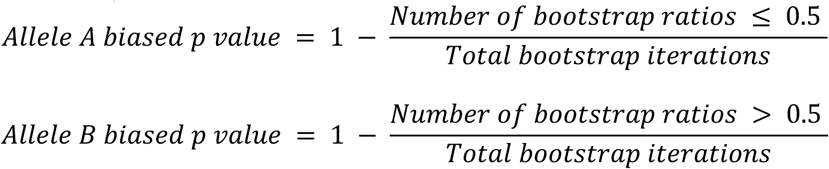

To identify ASE, we calculated adjusted p-values following multiple testing corrections. Because empirical p-values were derived from one-sided tests, they were converted to two-sided values by multiplication by two. Zero p-values were replaced with 1×10^−5^, corresponding to the resolution determined by 100,000 bootstrap iterations, prior to correction. Multiple testing correction was then performed using the Holm–Bonferroni method (68). Finally, genes were classified as ASE when the adjusted p-value was less than 0.05.

For tissue datasets with only three biological replicates, we additionally performed a count-based statistical test of ASE calls. Genes showing complete allelic bias, defined as an allele-bias ratio of consistently 0 or 1 in all three biological replicates with the same direction of allelic bias, were excluded from the beta-binomial test because their ASE patterns were unambiguous and fully consistent across replicates. For the remaining genes, allele-specific read counts were analyzed using a beta-binomial model implemented in glmmTMB (69). The beta-binomial model accounts for differences in read depth as well as variation among biological replicates. For each gene, we tested whether the expected allelic ratio differed from 0.5, representing equal expression of the B6 and CAST alleles. The resulting P values were corrected for multiple testing using the Benjamini–Hochberg procedure (70), and genes with an FDR < 0.05 were considered statistically supported. Among the genes identified as ASE (n = 252, 238, 317) by the bootstrap approach, this criterion excluded 30, 63, and 37 genes from beta-binomial testing in kidney, liver, and lung, respectively.

As a technical comparison of SNP-counting methods, we used an in-house custom Python script (count_snp.py) developed in a previous study (26).

### Measurement of mRNA expression and protein abundance across embryo development

RNA expression data were obtained from two previously published datasets (26,71). Transcriptomic data for the zygote through 8-cell stages were derived from Ozadam et al., whereas 16-cell and 32-cell stage data were obtained from Ghatpande et al. Gene expression levels were quantified using Reads Per Kilobase of transcript per Million mapped reads (RPKM), and RNA expression was compared between genes with and without ASE at each developmental stage. Mass spectrometry-based measurements of protein abundance were obtained (40).

### Quality assessment and correlation analysis

RNA-seq and ribosome profiling data quality was first assessed across all biological replicates. Spearman’s correlation coefficients were calculated using allele-specific read counts to assess correlations among biological replicates within each dataset (S1-3 Fig).

### TE calculation and analyses

After we selected highly correlated samples, we used two methods to calculate TE; compositional and ratio TE. For compositional TE calculation, we used the approach described previously (35). Prior to TE calculation, zero counts were imputed using multiplicative replacement via the zCompositions R package (v1.6.0; GBM algorithm). (72). RNA and ribosome profiling read counts were then normalized within each sample using the centered log-ratio transformation (73), and were converted to isometric log-ratio coordinates (74). A linear model was fitted between the two components, and the residuals from this model were defined as TE for each gene in a given sample.

Matched RNA-seq and ribosome profiling data were obtained from the same lysate for mESCs (13 clones), NPCs (5 clones), and adult tissues (n = 3 per organ). In contrast, embryo datasets were not matched (4 BxC and 3 CxB RNA-seq samples; 4 BxC and 4 CxB ribosome profiling samples). Therefore, TE was estimated by modeling all possible RNA–ribosome profiling combinations within each cross. Specifically, 16 BxC and 12 CxB RNA–Ribo pairs were generated and used for TE estimation using a compositional linear regression model. Then, average TE values were calculated across replicates for each gene and used for subsequent analyses.

As an alternative to compositional TE, we also calculated a conventional ratio-based measure of TE. For each gene, TE was defined as the ratio of ribosome occupancy to RNA abundance.

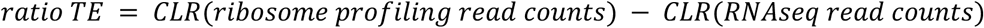

Prior to comparing TE (both compositional and ratio TE) between genes with and without ASE, we matched the two groups by RNA expression level to minimize potential effects of differences in transcript abundance. RNA expression was normalized using the centered log-ratio (CLR) transformation, and X-chromosome genes were excluded. Matching was performed in R using MatchIt version 4.5.3 (75). Because RNA expression was the sole matching covariate, genes with and without ASE were matched directly on CLR-normalized RNA expression using greedy nearest-neighbor matching. Balance in RNA expression between the matched groups was assessed using the standardized mean difference. TE values averaged across replicates were subsequently compared between the matched groups.

For statistical comparisons of averaged TE between genes with and without ASE, both compositional and ratio TE estimates were analyzed. TE differences between genes with and without ASE were assessed using two-sided Wilcoxon rank-sum tests. The effect size was defined as the difference in mean TE between the two groups and calculated as

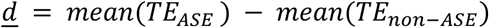

95 percent confidence intervals for the mean difference were estimated by nonparametric bootstrap resampling (N=5,000) with replacement.

### Allelic read count heterogeneity analysis

To assess heterogeneity in allelic bias across embryos or clones, we compared the allele-bias ratios at the gene level in each embryo or clone. For each gene, we calculated the mean allele-bias ratio across and the standard deviation as a measure of inter-clonal variability. These values were then visualized to compare both the average allelic bias and the degree of heterogeneity among clones across genes.

### Clonal allele-bias ratio and TE analysis

Using TE values calculated from the compositional regression model in matched RNA and ribosome profiling read counts from the same mESC clones, we performed clone-specific analyses. In each clone, paired TE and RNA abundance were obtained. For each clone, gene-level TE and RNA abundance estimates were obtained. To assess the relationship between clonal ASE and TE, TE values were compared between genes with ASE and genes without ASE after matching for RNA expression within each clone.

We further classified genes with ASE as showing either consistent or variable ASE based on their allele-bias ratios across individual clones. For this analysis, allele-bias ratios were first determined for each gene in each clone. Genes previously identified as showing ASE using our bootstrap-based method were then classified into two groups. A gene was classified as having consistent ASE if allele-bias ratio estimates were available in more than five informative clones and the allele-bias ratio was <0.2 or >0.8 in all informative clones. In contrast, a gene was classified as having variable ASE if allele-bias ratios between 0.2 and 0.8 were observed in more than one informative clone. These allele-bias ratio thresholds were adopted from Castel et al. (2015) to identify robust ASE while accounting for technical variability in allelic read counts arising from sequencing, mapping bias, and sampling error. TE was then compared between genes with consistent and variable ASE after matching for RNA expression.

As an additional analysis, we examined genes classified as without ASE by the bootstrap-based method that nevertheless showed allelic imbalance in individual clones. Within each clone, the TE of these genes was compared with that of genes showing allelic balance in the same clone after matching for RNA expression.

To assess whether the magnitude of allelic imbalance was associated with TE reduction, we further classified genes with ASE according to their clonal allele-bias ratios into three groups representing increasing degrees of allelic imbalance: 0.2 < ratio < 0.3 or 0.7 < ratio < 0.8; 0.1 < ratio < 0.2 or 0.8 < ratio < 0.9; and ratio < 0.1 or ratio > 0.9. TE was then compared across these groups. Genes without ASE in all clones were used as a reference group and were matched for RNA expression based on the average RNA expression of genes with ASE. P-values were calculated using the Wilcoxon rank-sum test by comparing each allele-bias ratio group with the genes without ASE group.

### ASE identification using independent RNA-seq datasets

To evaluate whether the observed TE patterns were consistent across independent datasets, we compared our results with published RNA-seq data. For embryos, we used 11 reciprocal 8-cell RNA-seq data from the Heard lab (43). Reads were trimmed using cutadapt with the following parameters:-m 20 -j 4 -a AGATCGGAAGAGCACACGTCTGAACTCCAGTCA -O 6.

For mESCs, we used RNA-seq data from single clones generated by the Spector Lab (41,43). We used 7 clone data for the analysis. Reads were trimmed using the following parameters:-m 20 -a AGATCGGAAGAGCACACGTCTGAACTCCAGTCA -A AGATCGGAAGAGCGTCGTGTAGGGAAAGAGTGT -O 6.

Subsequent processing, including SNP calling and ASE determination, was performed using the same pipeline described in the previous section to ensure consistency across datasets.

Because the published datasets did not include ribosome profiling data, TE could not be directly calculated from those studies. Therefore, after defining ASE using the external RNA-seq datasets, TE comparisons between genes with and without ASE were performed using TE values derived from our own matched RNA-seq and ribosome profiling datasets.

## DATA ACCESS

All data are available on Gene Expression Omnibus (embryo, mESC and NPC data, GSE343009, GSE343011; adult tissue data, GSE343750, GSE343578). All custom scripts used to perform bioinformatics analyses are available on GitHub: https://github.com/DayeaPark/Allele-specific-expression-and-translation

## ACKNOWLEDGMENTS

We thank Dr. David Spector for kindly providing hybrid mESCs. This work was supported by the National Institutes of Health grants (R35GM150667), as well as Welch Foundation grants (F-2027-20230405 and F-2027-20260402) (C.C.). D.G. acknowledges funding by the University of Lausanne and the Swiss National Science Foundation (grant 10002692 and NCCR RNA & Disease 205601), and bioinformatics and logistics support by Dr. A. Bulak Arpat and Dr. Virginie Ricci. We thank Dr. Jeannie Lee and Dr. Eric Aeby for collecting and providing the adult mouse tissues used to generate the data. We are grateful to Dr. Vighnesh Ghatpande and Dr. Shilpa Rao for insightful discussions that helped shape this work. We also thank all members of the Cenik laboratory for critical feedback on the manuscript. All original text in this paper was authored by the researchers. We acknowledge the assistance of Large Language Models for suggesting edits aimed at improving clarity and grammar, as well as in providing coding assistance during the development of this work. Some illustrations in this manuscript were created using BioRender (YC2A2CEEIH).

## AUTHOR CONTRIBUTIONS

D.P. conceived the study, performed experiments, analyzed the data, and wrote the manuscript.

C.C. supervised the study, provided guidance throughout the project, and edited the manuscript.

A.L. and D.G. generated the adult-tissue ribosome profiling and RNA-seq data and wrote the corresponding Methods section. All authors reviewed and approved the final manuscript.

## Supporting information

**S1 Figure.**
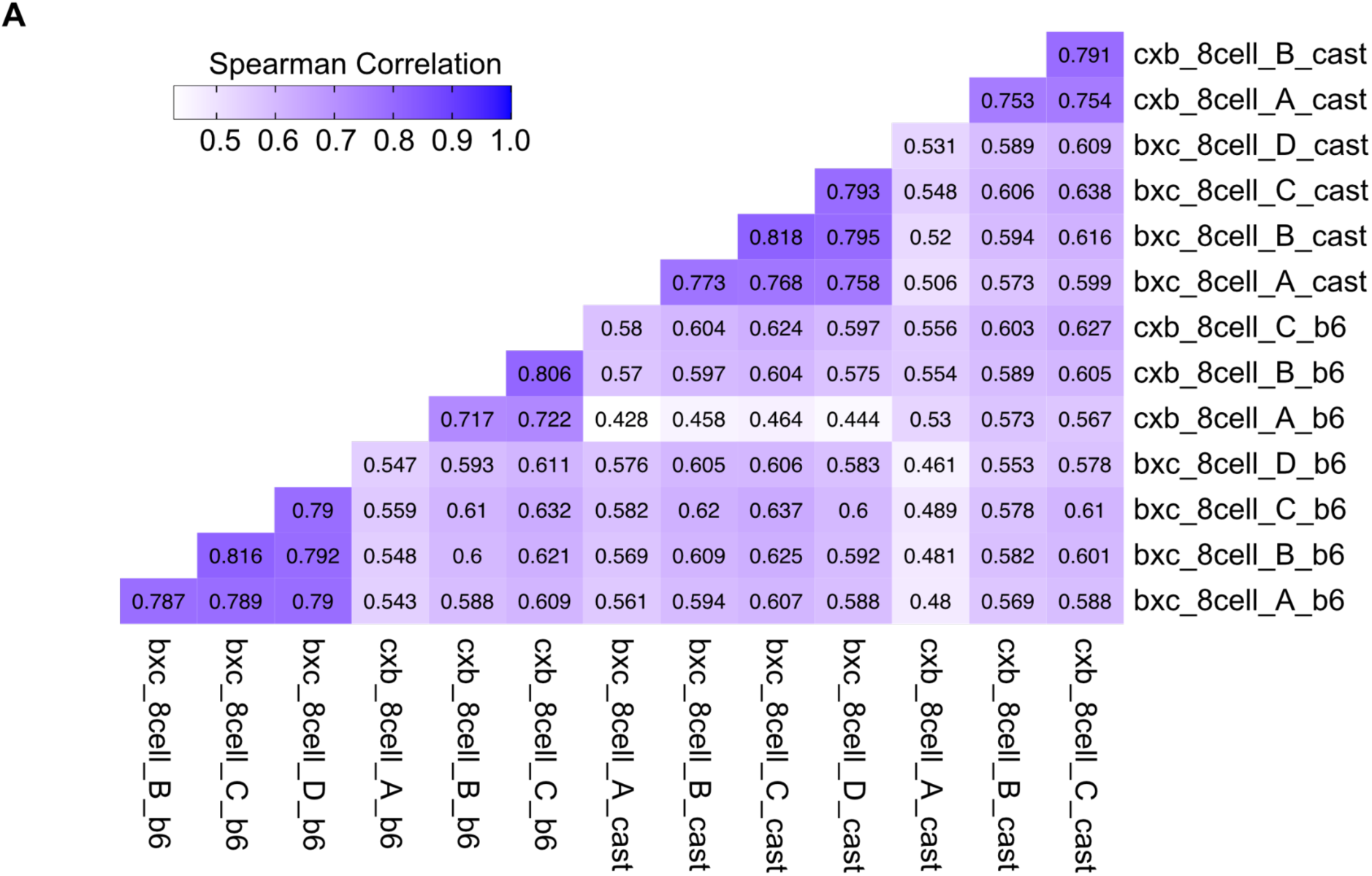
Correlations of allele-specific read counts of 8-cell reciprocal cross embryo. (A) Allelic read count correlations. Spearman’s correlation was calculated between reciprocal crosses using gene-level allelic RNA-seq read counts. Allelic read counts were first assigned to individual SNPs and then summed across all informative SNPs within each gene separately for the B6 and CAST alleles.

**S2 Figure.**
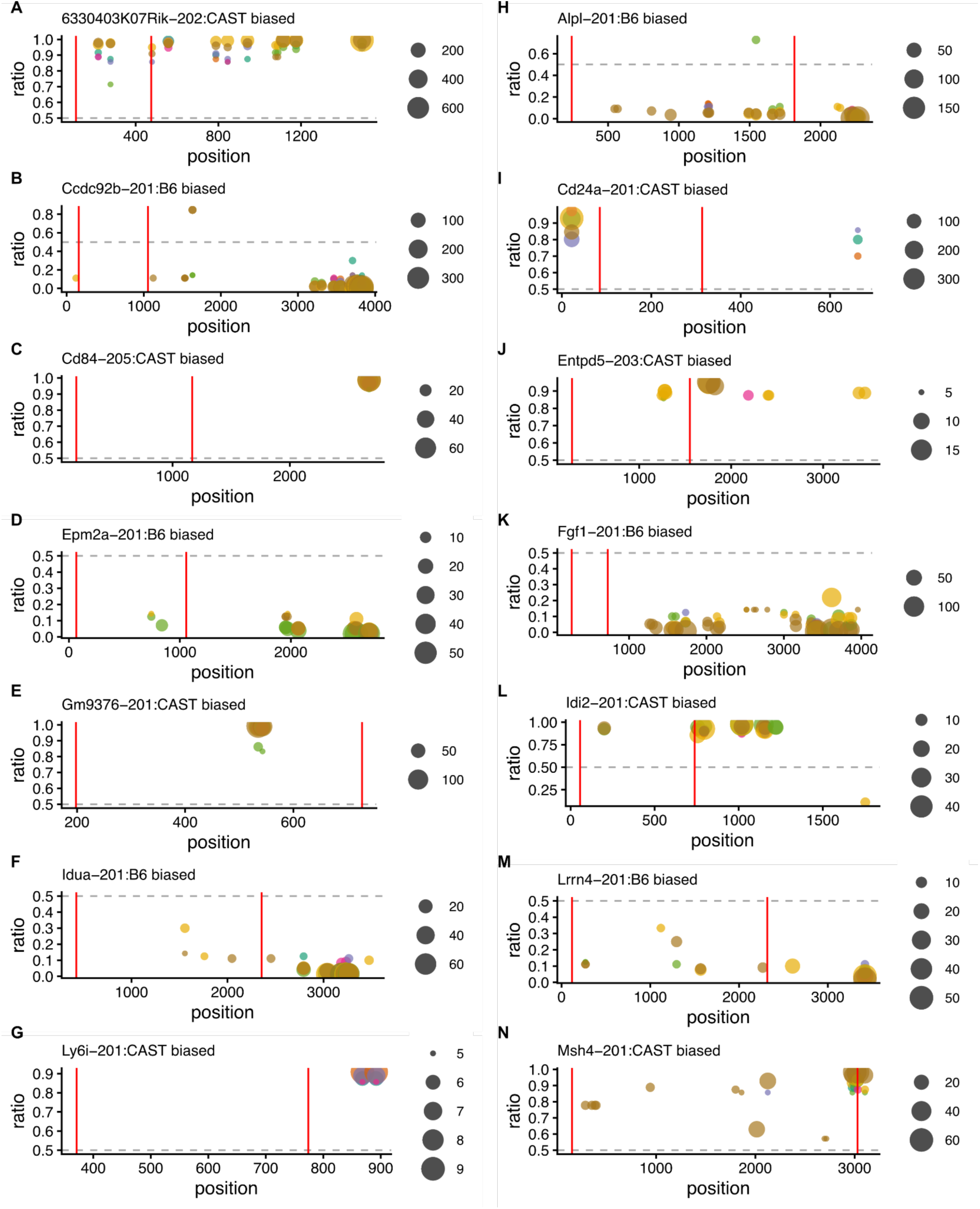
SNP-level allelic read support across representative genes. SNP-level allelic ratios and read coverage across representative genes. The y-axis shows the allele ratio at each informative SNP, and the x-axis indicates SNP position within the gene. Each color represents a distinct reciprocal 8-cell embryo, and dot size is proportional to the number of reads supporting each SNP. Red vertical lines indicate the boundaries between the 5′ UTR, coding sequence (CDS), and 3′ UTR.

**S3 Figure.**
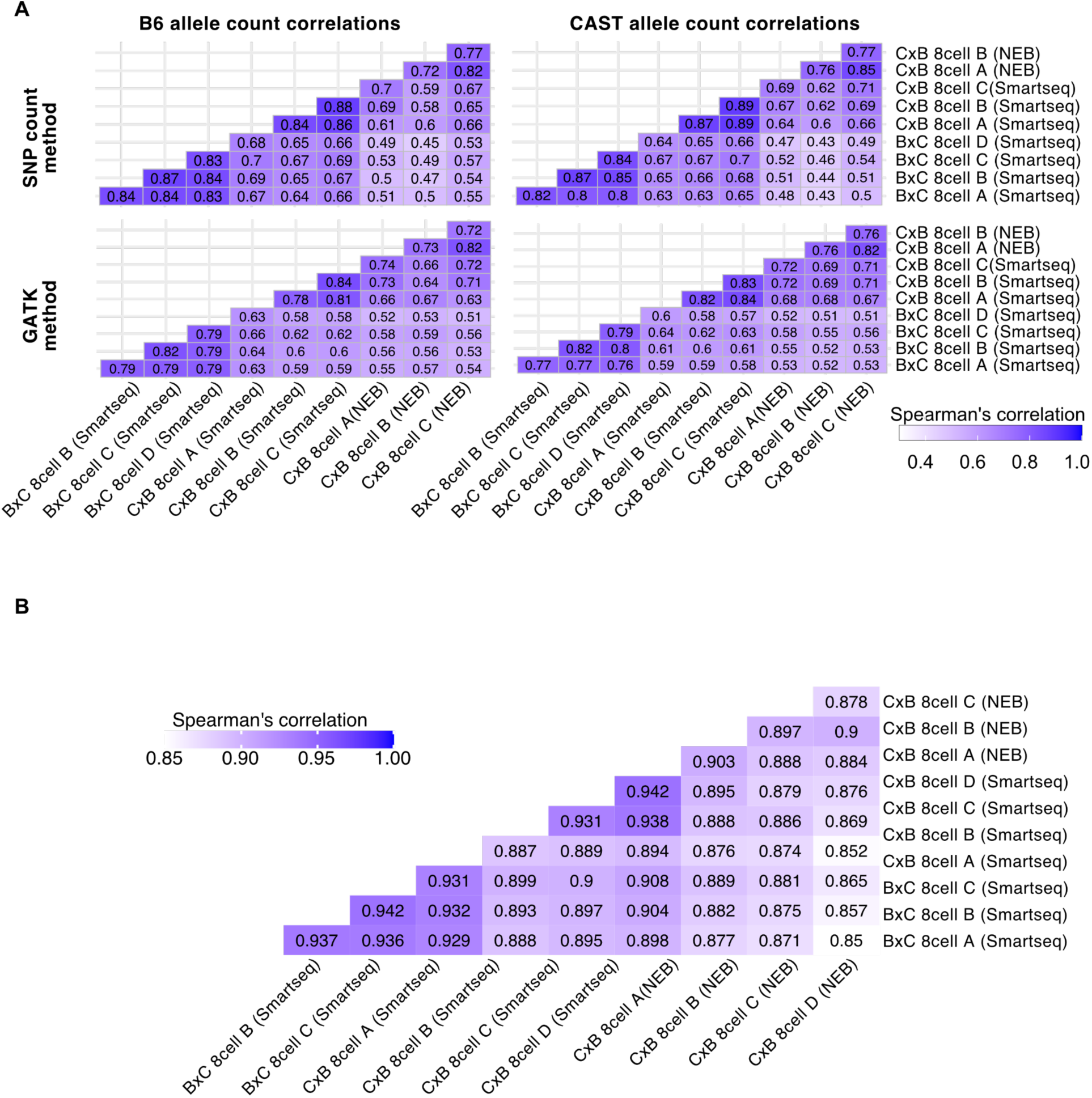
Sample correlations across computational and library preparation methods. (A) Correlations between B6 and CAST allele counts obtained using two SNP-calling methods: an in-house custom Python script called SNP count (top) and GATK (bottom). The left panels represent correlations for B6 alleles, and the right panels show correlations for CAST alleles. (B) RNA-seq read count correlations assessed using Spearman’s correlation coefficient.

**S4 Figure.**
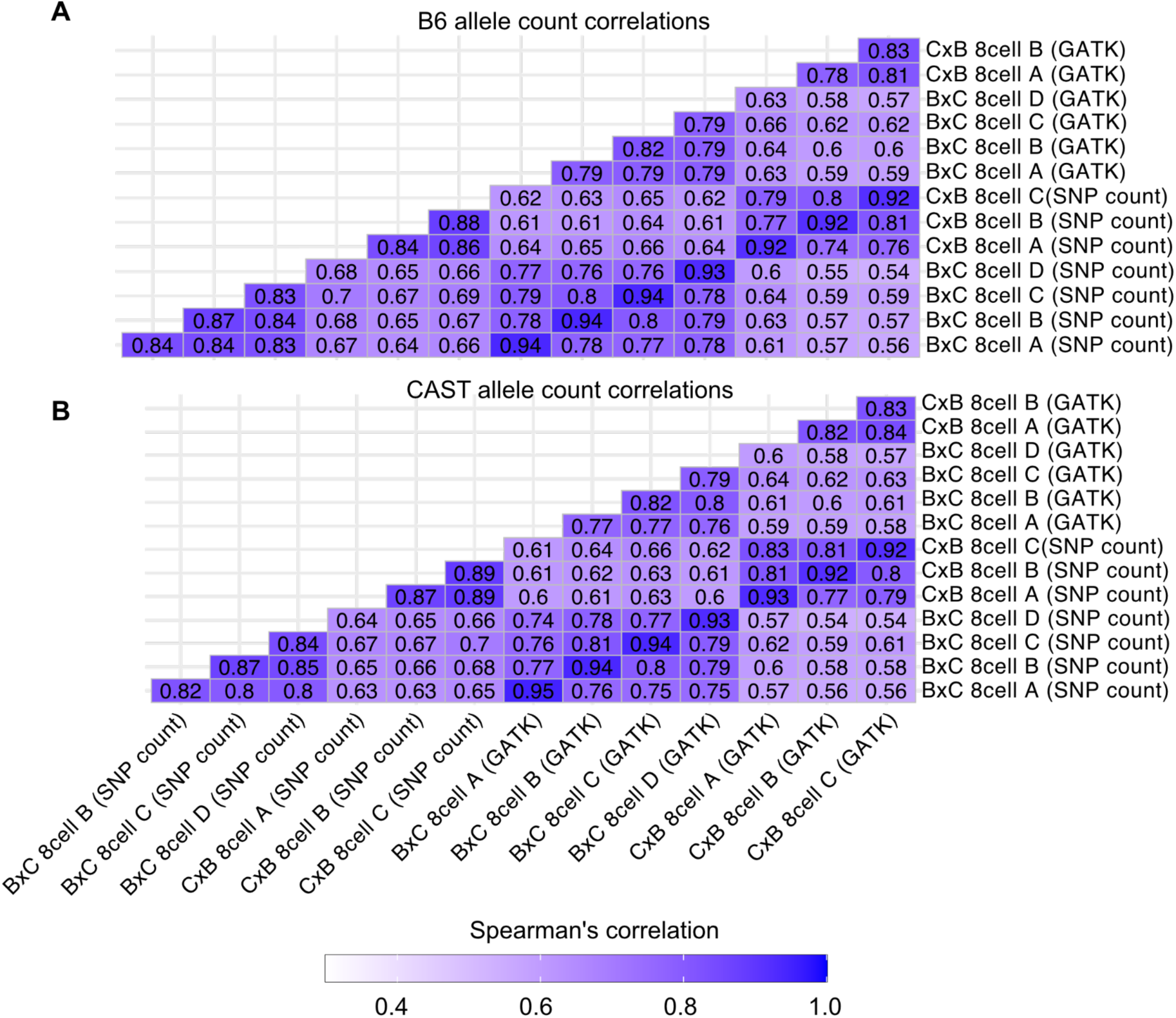
Allele read count correlations among different library preparation and SNP-calling method. Correlations between B6 (A) and CAST (B) allele counts obtained using two SNP-calling methods: an in-house custom Python script called SNP count and GATK, across embryo libraries generated by Smart-seq3.

**S5 Figure.**
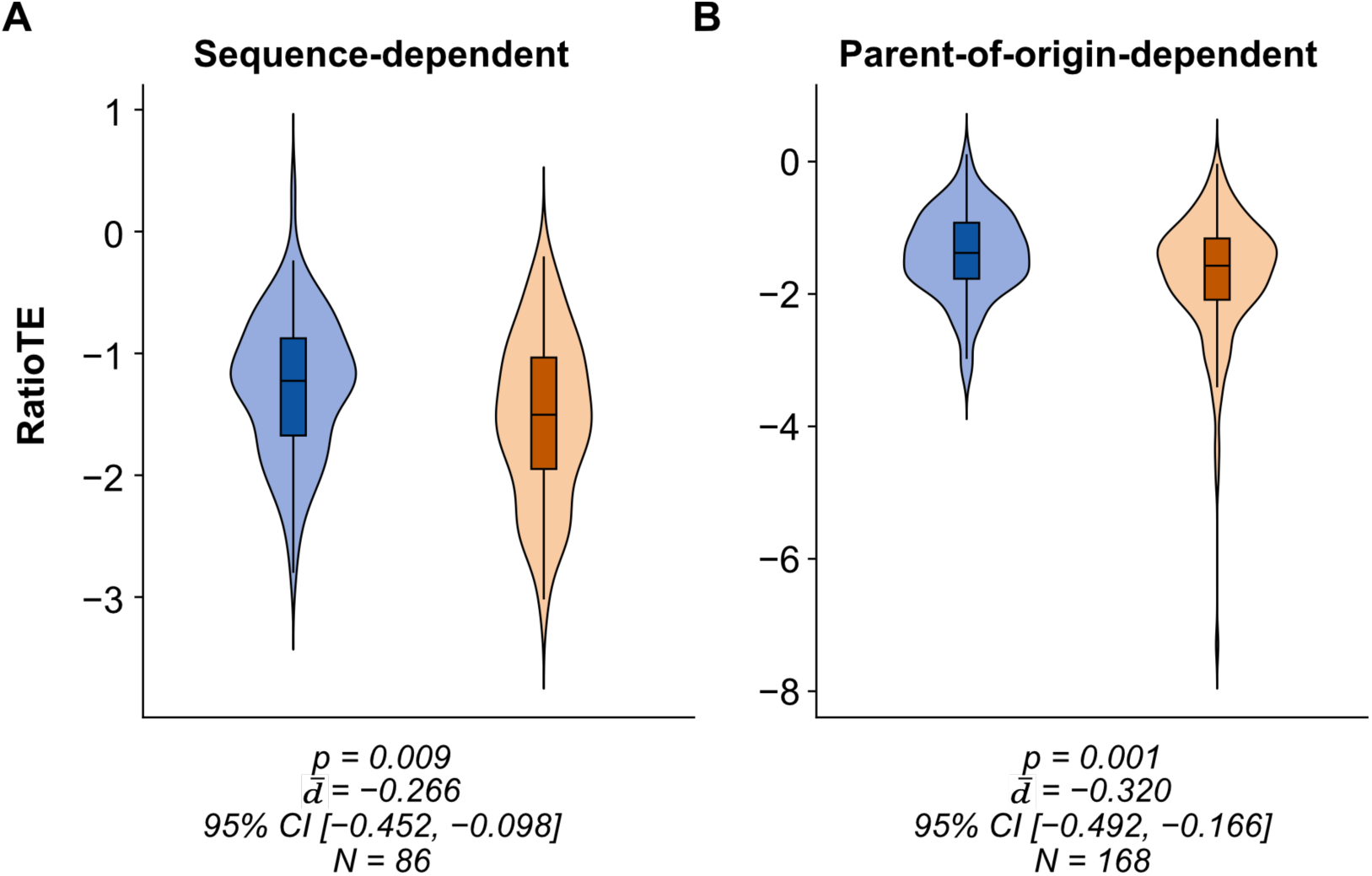
Ratio TE comparison in sequence-and parent-of-origin-dependent ASE. Ratio TE was calculated for 8-cell embryos and compared between genes with sequence-dependent ASE and non-ASE (A), and between genes with parent-of-origin-dependent ASE and non-ASE (B).

**S6 Figure.**
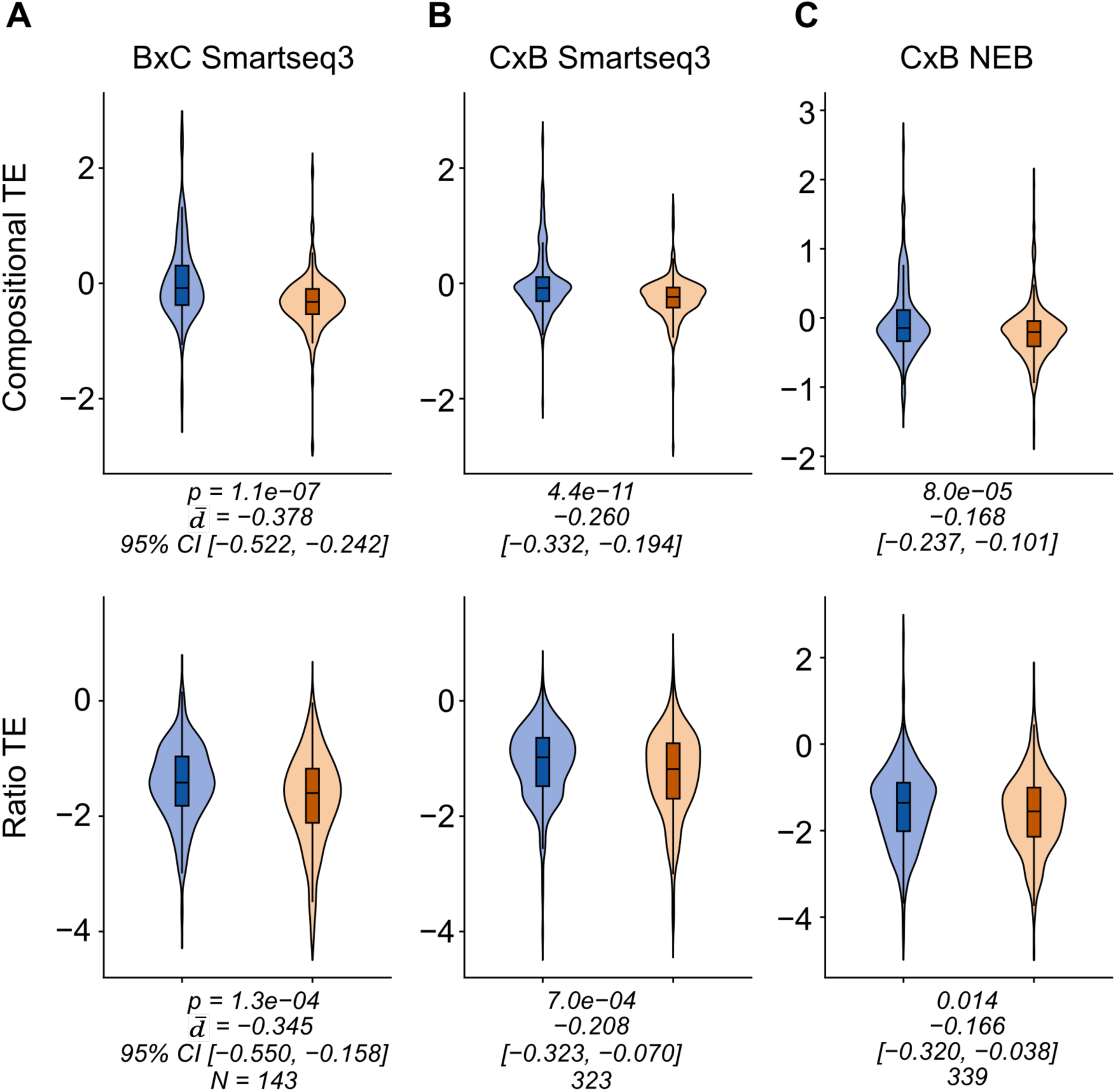
TE is consistent across library preparation methods. (A-C) TE measured in genes with ASE, as defined by GATK and a bootstrap method. The top panel shows compositional TE, while the bottom panels show the corresponding comparison using ratio TE. The genes with ASE identified in BxC 8-cell embryos generated using Smart-seq3 (A), CxB 8-cell generated using Smart-seq3 (B), and CxB 8-cell samples generated using NEB method (C). TE was compared between genes with and without ASE by matching from RNA expression.

**S7 Figure.**
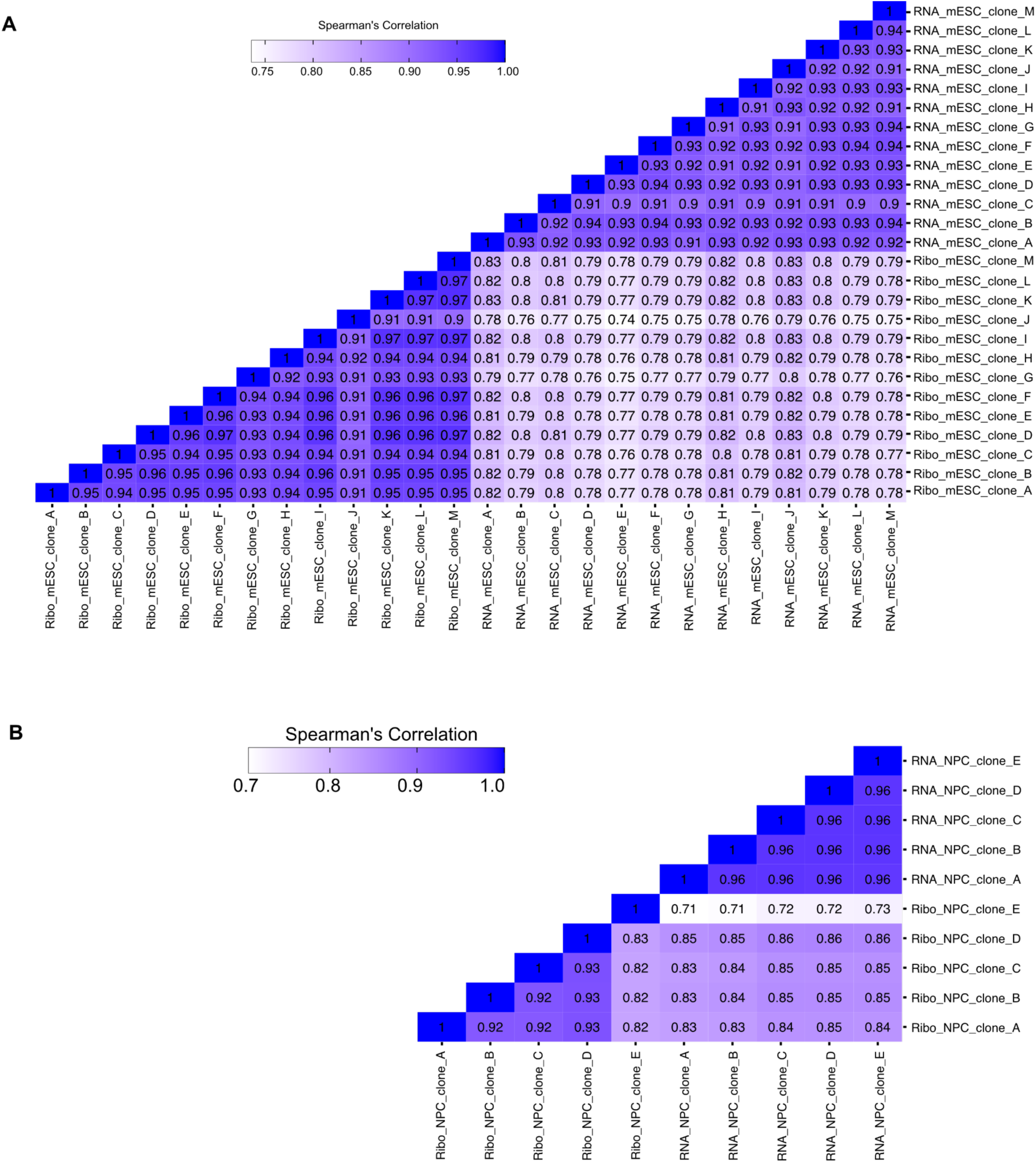
mESC clonal data quality control by measuring correlations. (A-B) RNA-seq and ribosome profiling sample correlations among mESC (A) and NPC (B) clones. Gene counts from RNA-seq and ribosome profiling data were generated using featureCounts and normalized using a centered log-ratio transformation. Spearman’s correlation was calculated using the normalized gene counts.

**S8 Figure.**
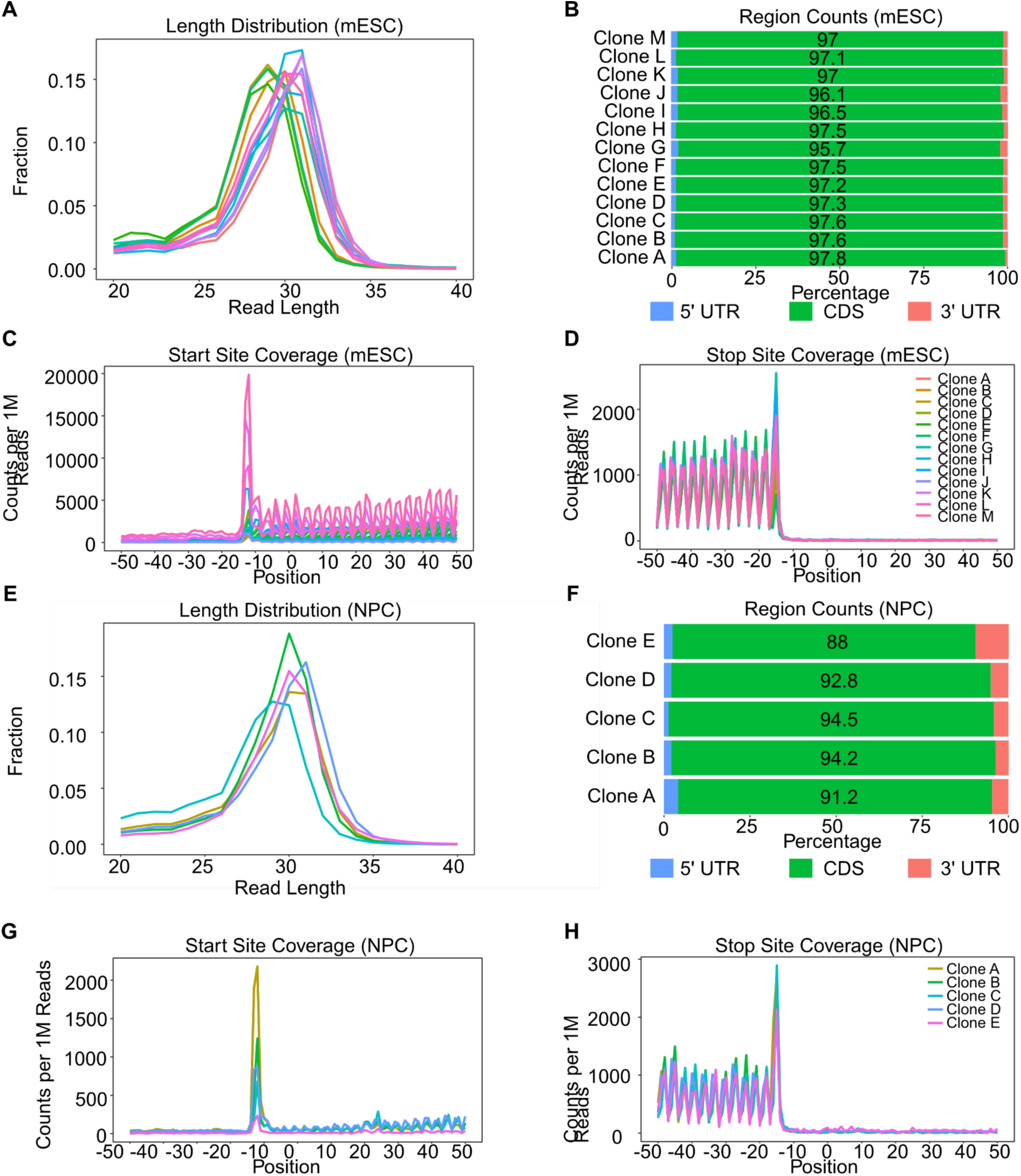
Quality control of mESC & NPC ribosome profiling data. (A) mESC ribosome profiling read length distribution. (B) Distribution of mESC ribosome profiling reads across genomic regions. (C–D) Metagene profiles of mESC ribosome profiling reads showing enrichment at annotated start codons (C) and stop codons (D). (E) NPC ribosome profiling read length distribution. (F) Distribution of NPC ribosome profiling reads across genomic regions. (G–H) Metagene profiles of NPC ribosome profiling reads showing enrichment at annotated start codons (G) and stop codons (H).\

**S9 Figure.**
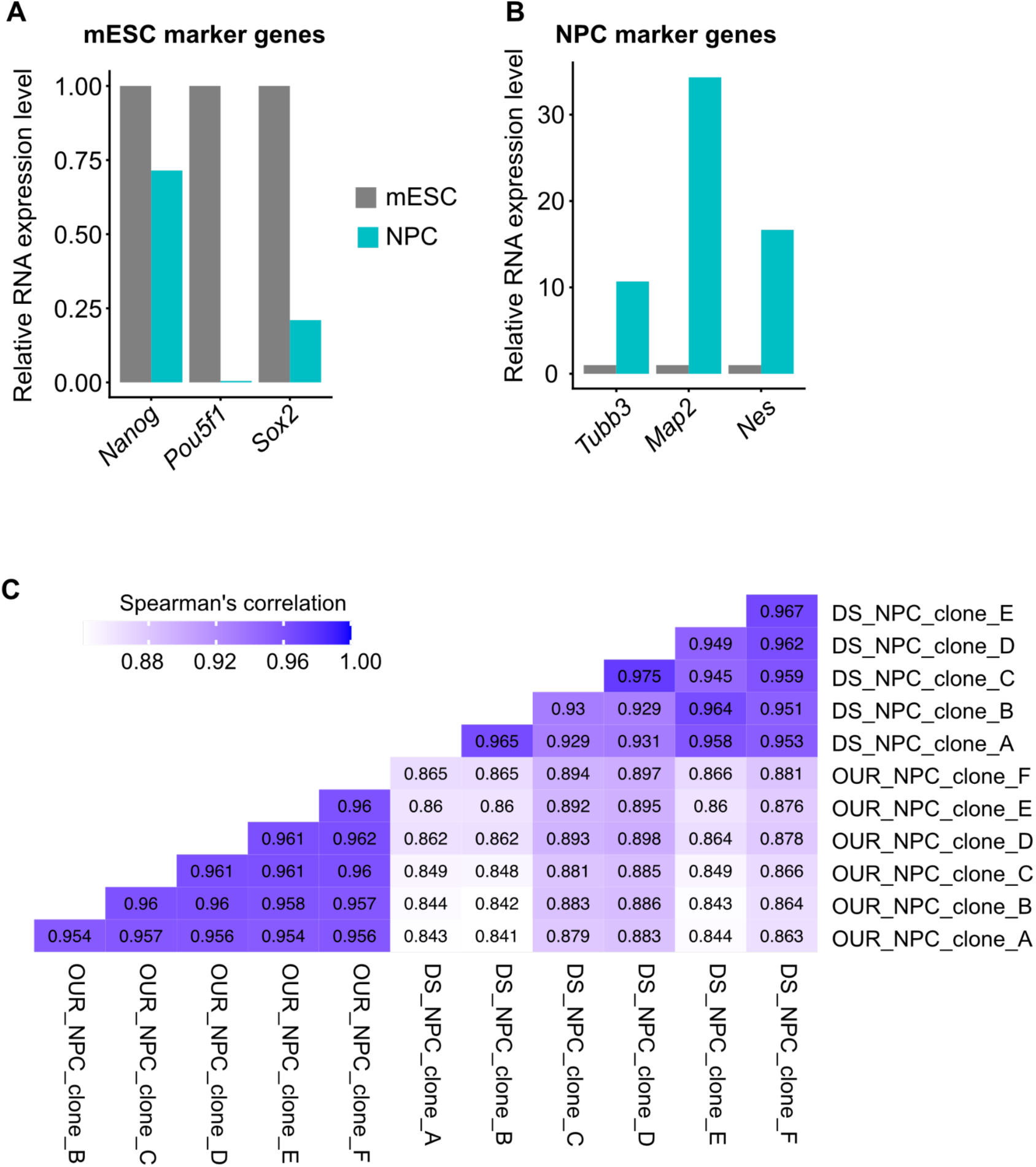
Validation of mESC differentiation into NPCs. (A-B) Relative expression of pluripotency and NPC marker genes measured by quantitative real-time PCR (qRT-PCR). Expression levels were normalized to Actb and calculated using the 2^−ΔΔCt method, confirming successful differentiation of mESCs into NPCs. (A) Expression of pluripotency marker genes (*Nanog*, *Pou5f1*[*Oct4*], and *Sox2*) (B) NPC marker genes (*Tubb3* [beta III-tubulin], Map2, and Nestin) measured by qRT-PCR. (C) Spearman correlation of gene expression (RNA-seq counts) between NPCs generated in this study (OUR) and a previously published NPC dataset (DS) (Eckersley-Maslin et al., 2014), demonstrating high transcriptomic similarity between the two datasets. OUR and DS correspond to the x-and y-axis labels, respectively.

**S10 Figure.**
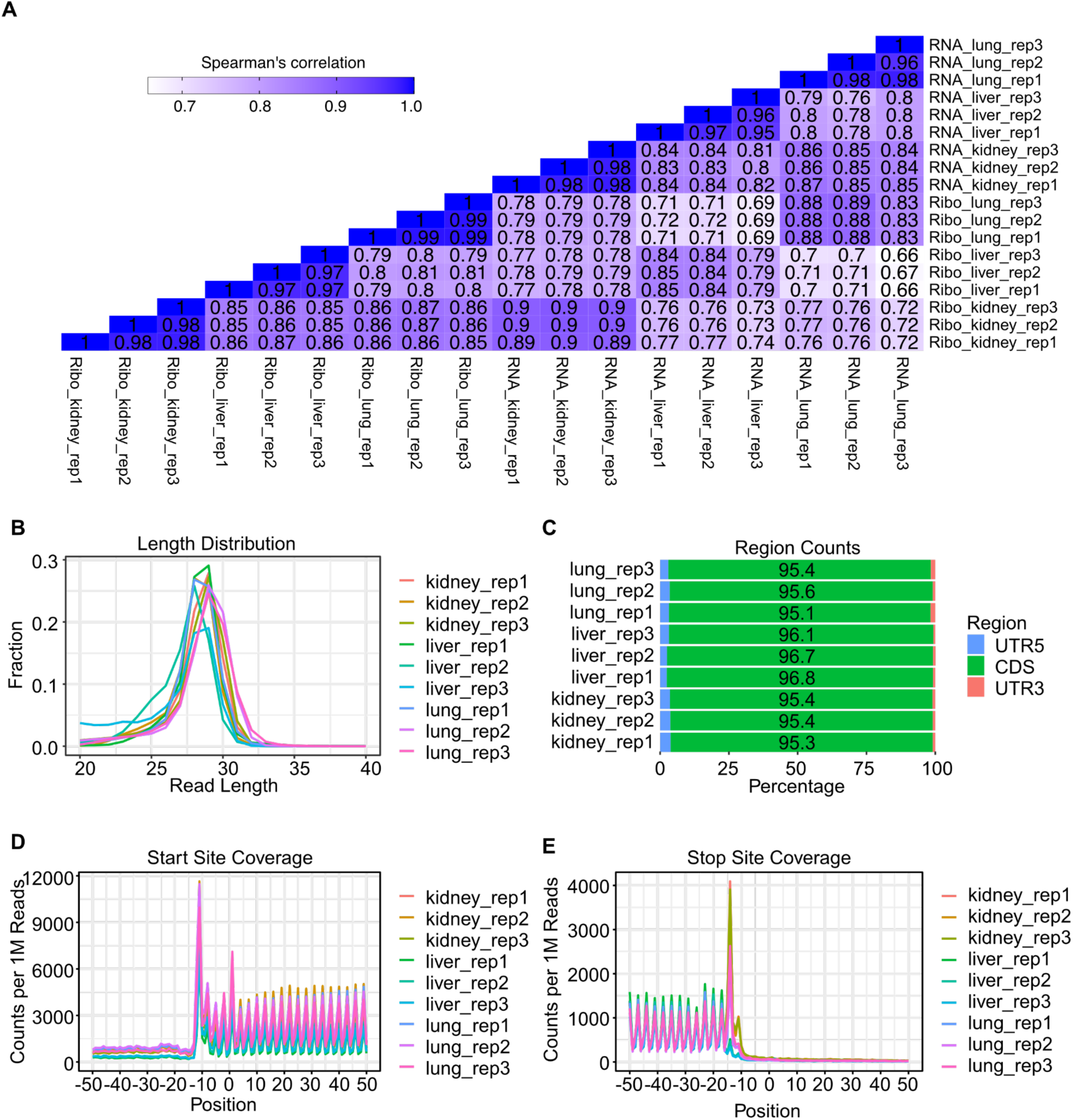
Data quality control for ribosome profiling and RNA-seq from adult tissues. (A) Spearman’s correlations between ribosome profiling and RNA-seq data across adult tissue samples. (B–E) Quality assessment of ribosome profiling data. (B) Distribution of ribosome profiling read lengths. (C) Distribution of ribosome profiling reads across genomic regions. (D–E) Metagene profiles of ribosome profiling reads showing enrichment around annotated start codons (D) and stop codons (E).

**S11 Figure.**
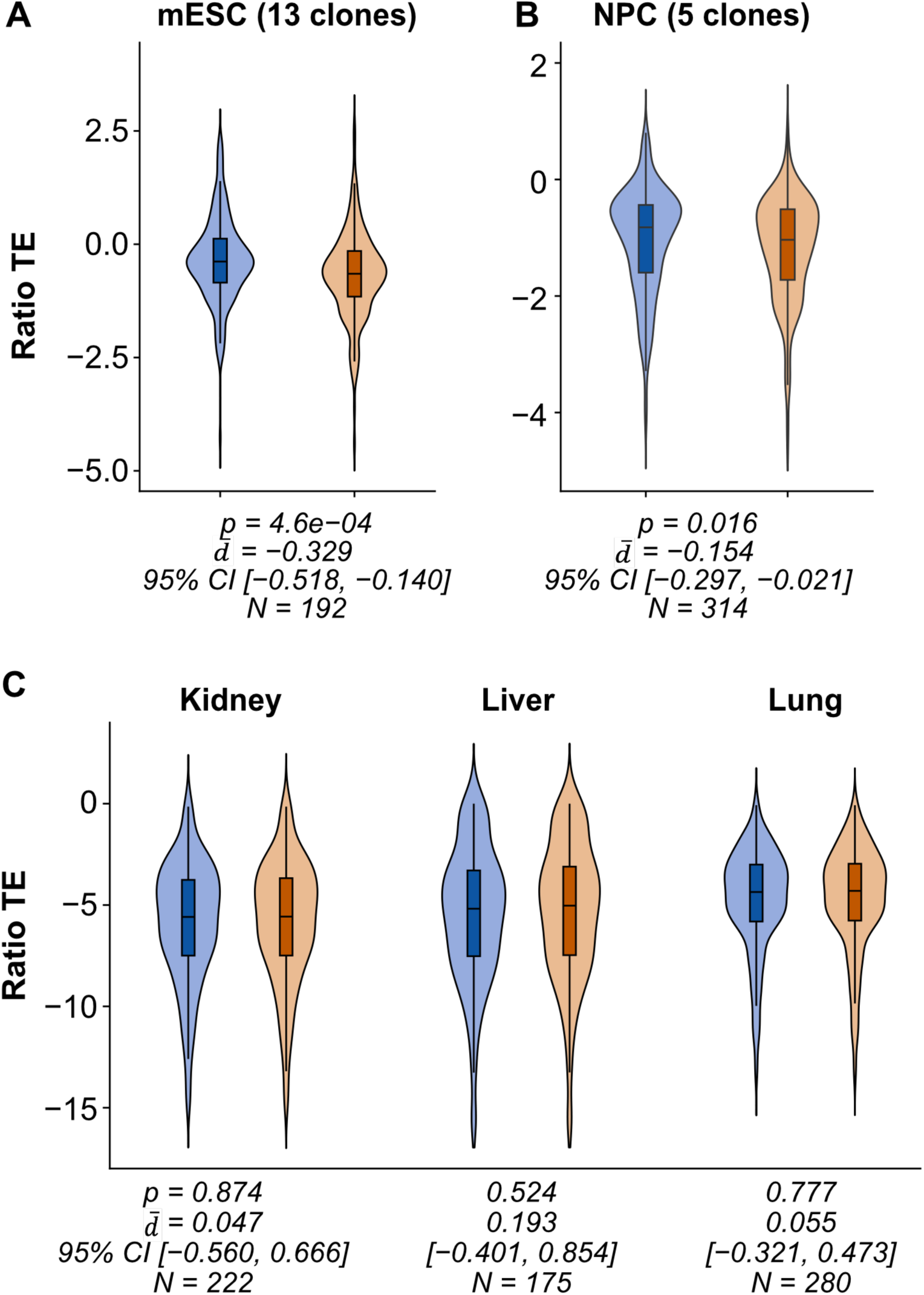
Ratio TE comparison among different cell types. Ratio TE was quantified in mESCs (A), NPCs (B), and adult tissues (C). The TE was compared between genes with and without ASE in each cell type.

**S12 Table.** Variable number of genes with ASE across clones.

| Clone name | genes with ASE | genes with consistent ASE | genes with variable ASE |
| --- | --- | --- | --- |
| clone_A | 164 | 29 | 135 |
| clone_B | 170 | 30 | 140 |
| clone_C | 127 | 26 | 101 |
| clone_D | 141 | 30 | 111 |
| clone_E | 136 | 26 | 110 |
| clone_F | 168 | 29 | 139 |
| clone_G | 152 | 28 | 124 |
| clone_H | 158 | 28 | 130 |
| clone_I | 137 | 27 | 110 |
| clone_J | 176 | 29 | 147 |
| clone_K | 137 | 26 | 111 |
| clone_L | 148 | 28 | 120 |
| clone_M | 160 | 30 | 130 |

**S13 Table.** Statistical results of clone-specific TE comparisons.

| Clone name | Mean TE diff | Lower CI | Upper CI | P-value | Assigned figure |
| --- | --- | --- | --- | --- | --- |
| clone_A | -0.3061159 | -0.4545688 | -0.1645509 | 3.228E-05 | Fig 4B |
| clone_B | -0.3605781 | -0.5101841 | -0.2135987 | 2.422E-06 | Fig 4B |
| clone_C | -0.3953156 | -0.5723718 | -0.2324012 | 8.542E-05 | Fig 4B |
| clone_D | -0.4324598 | -0.5971937 | -0.2565586 | 1.635E-05 | Fig 4B |
| clone_E | -0.2829746 | -0.467558 | -0.1059 | 1.499E-03 | Fig 4B |
| clone_F | -0.4494516 | -0.6122326 | -0.2877455 | 7.659E-07 | Fig 4B |
| clone_G | -0.3393925 | -0.4852568 | -0.1998415 | 1.702E-06 | Fig 4B |
| clone_H | -0.3336358 | -0.4894212 | -0.1776312 | 1.057E-04 | Fig 4B |
| clone_I | -0.4008234 | -0.5830185 | -0.2231878 | 7.994E-05 | Fig 4B |
| clone_J | -0.2998893 | -0.4479359 | -0.1597237 | 5.250E-05 | Fig 4B |
| clone_K | -0.4352028 | -0.5930013 | -0.267252 | 6.907E-06 | Fig 4B |
| clone_L | -0.4084746 | -0.5780717 | -0.240532 | 8.267E-06 | Fig 4B |
| clone_M | -0.411257 | -0.569253 | -0.2423632 | 3.634E-06 | Fig 4B |
| clone_A | -0.4471421 | -0.683625 | -0.2053219 | 7.524E-04 | Fig 4D |
| clone_B | -0.4663384 | -0.7467611 | -0.1829918 | 8.707E-04 | Fig 4D |
| clone_C | -0.4084871 | -0.6392785 | -0.1666171 | 8.464E-03 | Fig 4D |
| clone_D | -0.42856 | -0.7231808 | -0.1122173 | 5.449E-03 | Fig 4D |
| clone_E | -0.3026948 | -0.6195446 | 0.02624381 | 5.128E-02 | Fig 4D |
| clone_F | -0.4153183 | -0.6898146 | -0.1075182 | 4.185E-03 | Fig 4D |
| clone_G | -0.4009483 | -0.6759685 | -0.0905761 | 1.361E-03 | Fig 4D |
| clone_H | -0.4662051 | -0.6918893 | -0.1917623 | 1.132E-03 | Fig 4D |
| clone_I | -0.4085073 | -0.7216843 | -0.0969302 | 9.078E-03 | Fig 4D |
| clone_J | -0.4423709 | -0.7024946 | -0.1980303 | 8.335E-04 | Fig 4D |
| clone_K | -0.5002278 | -0.7657372 | -0.1942194 | 7.584E-04 | Fig 4D |
| clone_L | -0.4347324 | -0.7037696 | -0.1202067 | 3.795E-03 | Fig 4D |
| clone_M | -0.4580293 | -0.748738 | -0.1701602 | 1.646E-03 | Fig 4D |
| clone_A | -0.1671896 | -0.3136235 | -0.0278919 | 2.268E-02 | Fig 4E |
| clone_B | -0.2172771 | -0.3716408 | -0.063309 | 4.098E-03 | Fig 4E |
| clone_C | -0.2559072 | -0.4346452 | -0.0927428 | 1.722E-02 | Fig 4E |
| clone_D | -0.2822727 | -0.4560228 | -0.101809 | 7.908E-03 | Fig 4E |
| clone_E | -0.1850652 | -0.3626559 | -0.0075372 | 3.882E-02 | Fig 4E |
| clone_F | -0.3282235 | -0.4792777 | -0.1731901 | 7.100E-04 | Fig 4E |
| clone_G | -0.2124909 | -0.36155 | -0.0644967 | 2.845E-03 | Fig 4E |
| clone_H | -0.1889746 | -0.3356971 | -0.0314101 | 3.951E-02 | Fig 4E |
| clone_I | -0.2660043 | -0.4379512 | -0.0748049 | 1.484E-02 | Fig 4E |
| clone_J | -0.1707544 | -0.3159931 | -0.0342082 | 2.530E-02 | Fig 4E |
| clone_K | -0.2731914 | -0.4310707 | -0.0926925 | 1.021E-02 | Fig 4E |
| clone_L | -0.2691759 | -0.4448672 | -0.1003961 | 4.893E-03 | Fig 4E |
| clone_M | -0.2649119 | -0.4174246 | -0.0934435 | 4.425E-03 | Fig 4E |
| clone_A | -0.1204653 | -0.2052144 | -0.0322977 | 2.431E-03 | Fig 4F |
| clone_B | -0.104439 | -0.2090963 | 0.00071002 | 3.149E-02 | Fig 4F |
| clone_C | -0.1310536 | -0.255988 | -0.0140167 | 8.312E-02 | Fig 4F |
| clone_D | -0.1861107 | -0.3162469 | -0.0612574 | 7.754E-03 | Fig 4F |
| clone_E | -0.0840341 | -0.1917758 | 0.03100695 | 1.027E-01 | Fig 4F |
| clone_F | -0.1483925 | -0.2578502 | -0.0445218 | 5.225E-04 | Fig 4F |
| clone_G | -0.1388723 | -0.2365531 | -0.033026 | 1.820E-03 | Fig 4F |
| clone_H | -0.1483495 | -0.2447558 | -0.0473424 | 3.475E-03 | Fig 4F |
| clone_I | -0.1570071 | -0.2692232 | -0.0508537 | 6.773E-03 | Fig 4F |
| clone_J | -0.2860657 | -0.3837484 | -0.1860205 | 7.295E-09 | Fig 4F |
| clone_K | -0.1171106 | -0.2430502 | -0.0017627 | 2.107E-02 | Fig 4F |
| clone_L | -0.2184121 | -0.3301483 | -0.0931717 | 1.746E-04 | Fig 4F |
| clone_M | -0.1346156 | -0.2451835 | -0.0149947 | 4.123E-03 | Fig 4F |
| clone_A | -0.3418432 | -0.5639161 | -0.1392208 | 1.071E-03 | S15A Fig |
| clone_B | -0.4178089 | -0.6354672 | -0.1888069 | 1.500E-04 | S15A Fig |
| clone_C | -0.3683255 | -0.6231703 | -0.1324389 | 5.076E-03 | S15A Fig |
| clone_D | -0.4627411 | -0.7204311 | -0.2265117 | 3.237E-04 | S15A Fig |
| clone_E | -0.2922686 | -0.5448176 | -0.0407799 | 5.695E-03 | S15A Fig |
| clone_F | -0.5634595 | -0.7782917 | -0.3401972 | 8.996E-07 | S15A Fig |
| clone_G | -0.3416735 | -0.5673076 | -0.1194284 | 2.326E-03 | S15A Fig |
| clone_H | -0.3691038 | -0.5739256 | -0.1459287 | 1.183E-03 | S15A Fig |
| clone_I | -0.4984798 | -0.7564173 | -0.2453968 | 3.296E-04 | S15A Fig |
| clone_J | -0.2755369 | -0.4800723 | -0.0697364 | 7.122E-03 | S15A Fig |
| clone_K | -0.5410029 | -0.7783093 | -0.3142685 | 4.268E-05 | S15A Fig |
| clone_L | -0.5031843 | -0.7306182 | -0.2768212 | 1.062E-04 | S15A Fig |
| clone_M | -0.4977733 | -0.7323049 | -0.2612629 | 6.419E-05 | S15A Fig |
| clone_A | -0.7083811 | -1.0284879 | -0.3674158 | 4.192E-04 | S15B Fig |
| clone_B | -0.7330429 | -1.1204501 | -0.3281076 | 9.207E-04 | S15B Fig |
| clone_C | -0.4899638 | -0.822621 | -0.1728978 | 3.930E-02 | S15B Fig |
| clone_D | -0.6138788 | -1.0376833 | -0.1713902 | 7.368E-03 | S15B Fig |
| clone_E | -0.5052303 | -0.9613888 | -0.0473482 | 2.666E-02 | S15B Fig |
| clone_F | -0.6540116 | -1.0340923 | -0.2431047 | 2.657E-03 | S15B Fig |
| clone_G | -0.6230975 | -1.0408971 | -0.1781401 | 2.642E-03 | S15B Fig |
| clone_H | -0.724003 | -1.0908796 | -0.342911 | 1.260E-03 | S15B Fig |
| clone_I | -0.6103083 | -1.0886732 | -0.1920662 | 1.357E-02 | S15B Fig |
| clone_J | -0.7751408 | -1.1649789 | -0.4060876 | 2.848E-04 | S15B Fig |
| clone_K | -0.7818876 | -1.1560282 | -0.363421 | 5.921E-04 | S15B Fig |
| clone_L | -0.6723027 | -1.084207 | -0.2074278 | 5.078E-03 | S15B Fig |
| clone_M | -0.7234143 | -1.1756867 | -0.2860915 | 1.817E-03 | S15B Fig |
| clone_A | -0.1171307 | -0.3096515 | 0.08138459 | 1.978E-01 | S15C Fig |
| clone_B | -0.1877636 | -0.4069498 | 0.03144524 | 5.224E-02 | S15C Fig |
| clone_C | -0.1964808 | -0.4255415 | 0.03444665 | 1.129E-01 | S15C Fig |
| clone_D | -0.2401787 | -0.5013188 | 0.00294626 | 4.683E-02 | S15C Fig |
| clone_E | -0.1220172 | -0.3594674 | 0.12962702 | 1.367E-01 | S15C Fig |
| clone_F | -0.3681268 | -0.5959353 | -0.1542763 | 1.063E-03 | S15C Fig |
| clone_G | -0.1379358 | -0.3611586 | 0.09628987 | 1.841E-01 | S15C Fig |
| clone_H | -0.1396173 | -0.3665277 | 0.09068737 | 1.521E-01 | S15C Fig |
| clone_I | -0.2930056 | -0.5266272 | -0.0157452 | 3.069E-02 | S15C Fig |
| clone_J | -0.0410466 | -0.2525484 | 0.15727082 | 4.842E-01 | S15C Fig |
| clone_K | -0.2825663 | -0.5179801 | -0.0376668 | 3.428E-02 | S15C Fig |
| clone_L | -0.2816806 | -0.52342 | -0.0358593 | 2.310E-02 | S15C Fig |
| clone_M | -0.2611049 | -0.4941102 | -0.0322549 | 2.659E-02 | S15C Fig |
| clone_A | -0.1407322 | -0.2580885 | -0.0177434 | 3.220E-02 | S15D Fig |
| clone_B | -0.1051034 | -0.2544772 | 0.04216118 | 7.890E-02 | S15D Fig |
| clone_C | -0.131104 | -0.2775705 | 0.02643444 | 1.441E-01 | S15D Fig |
| clone_D | -0.2541041 | -0.4197008 | -0.0859146 | 4.892E-03 | S15D Fig |
| clone_E | -0.0808761 | -0.2183912 | 0.05566772 | 3.024E-01 | S15D Fig |
| clone_F | -0.2046951 | -0.35621 | -0.0633562 | 2.439E-03 | S15D Fig |
| clone_G | -0.1561472 | -0.3114561 | -0.01688 | 1.800E-02 | S15D Fig |
| clone_H | -0.1688804 | -0.2932887 | -0.0363763 | 7.104E-03 | S15D Fig |
| clone_I | -0.1867637 | -0.3345878 | -0.040687 | 1.774E-02 | S15D Fig |
| clone_J | -0.2425691 | -0.3723153 | -0.1038673 | 4.233E-04 | S15D Fig |
| clone_K | -0.1759153 | -0.3327708 | -0.0297466 | 1.619E-02 | S15D Fig |
| clone_L | -0.2831858 | -0.4308481 | -0.1426689 | 4.870E-04 | S15D Fig |
| clone_M | -0.1812981 | -0.3413398 | -0.0159557 | 4.746E-03 | S15D Fig |

**S14 Figure.**
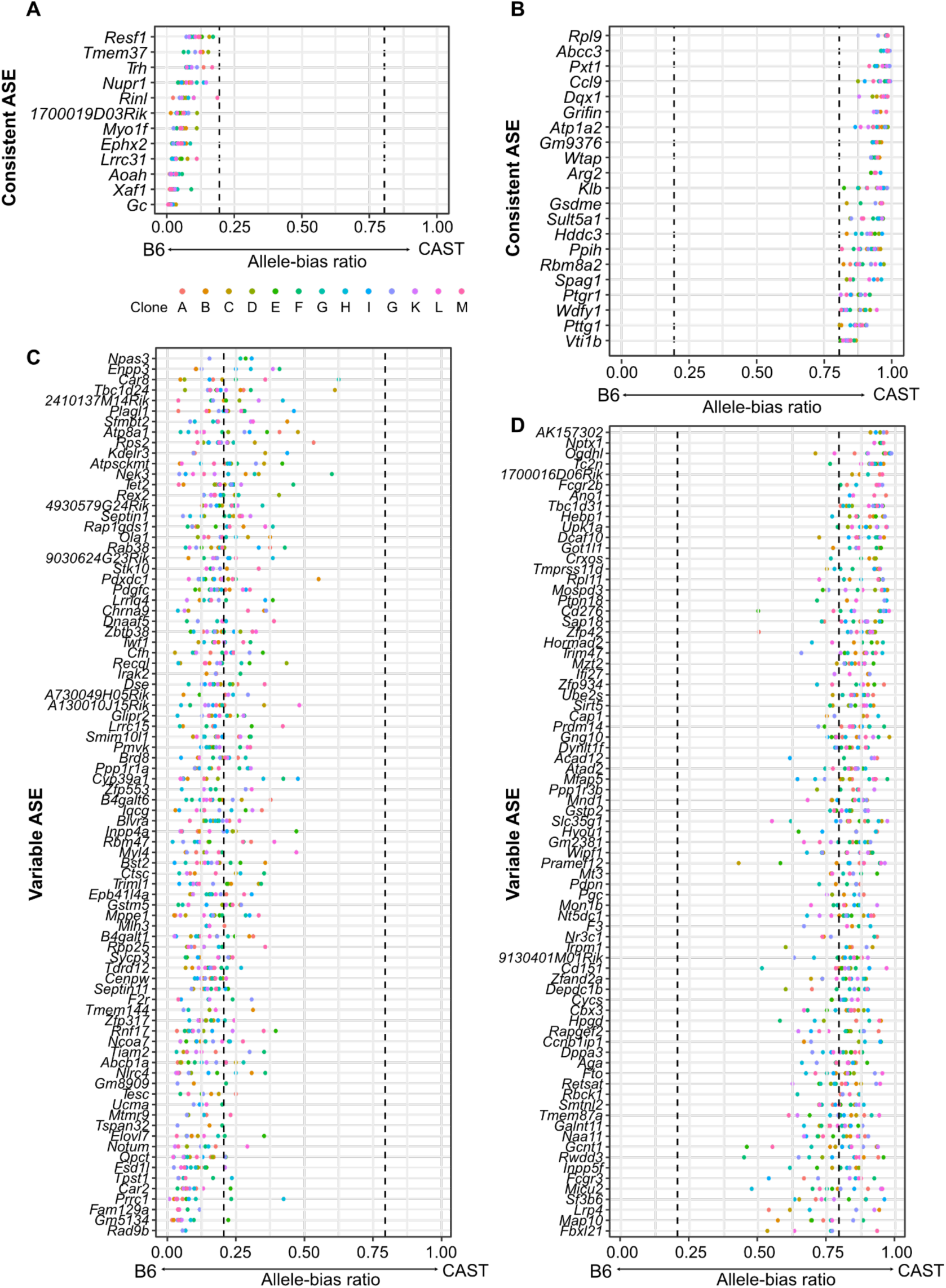
Classification of genes with consistent ASE and variable ASE in mESC. Clonal allele-bias ratio distributions used to classify genes as consistent ASE (A-B) or variable ASE (C-D). Genes were classified as consistent ASE if allele-bias ratios were available for more than five of the 13 mESC clones and all informative clones exhibited ASE in the same allelic direction (allele-bias ratio < 0.2 or > 0.8). Genes that did not meet these criteria were classified as variable ASE, including genes showing ASE in only a subset of informative clones, non-ASE in one or more clones, or allele-bias ratio estimates in five or fewer clones.

**S15 Figure.**
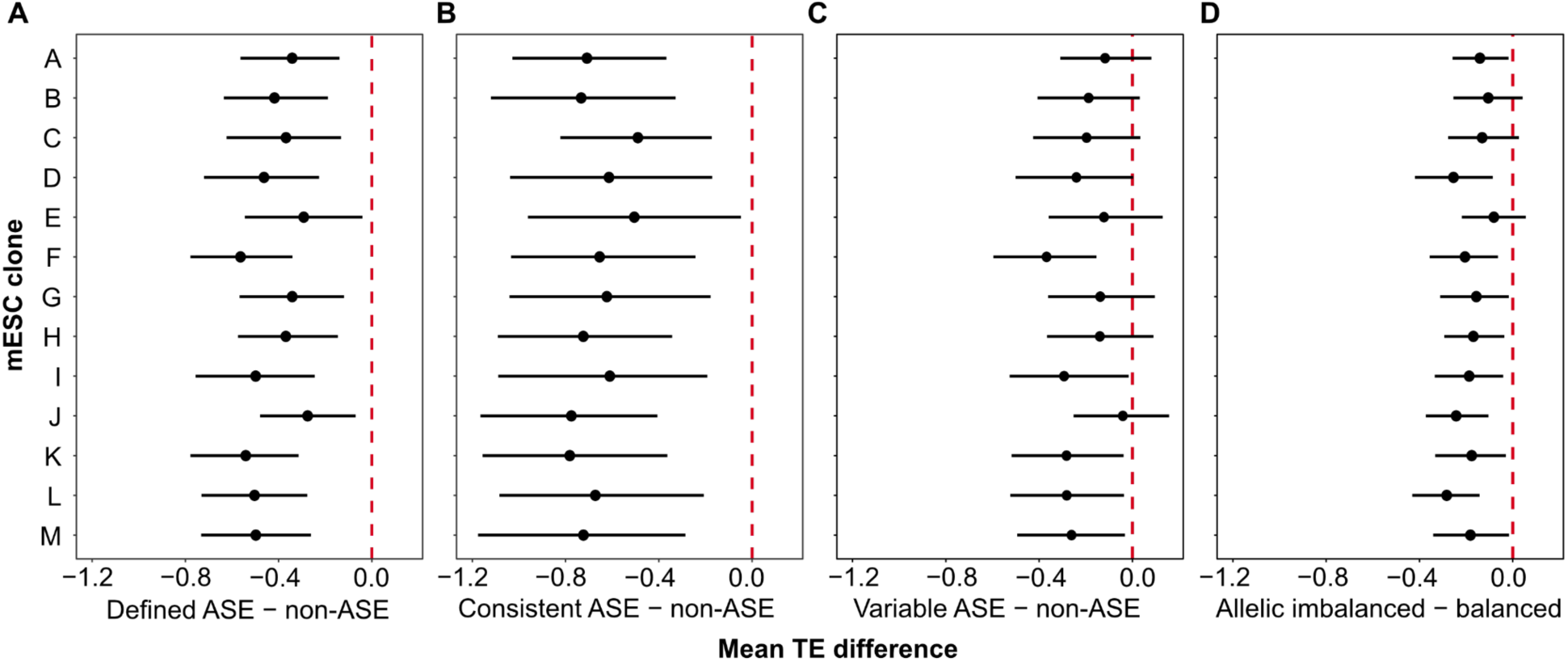
Clone-specific TE differences between genes with and without allelic imbalance using the ratio TE method. Clone-specific differences in ratio TE between genes with and without allelic bias, calculated as *TE*_’_*_with allice bias_*-*TE*_’_*_without allice bias_*. Comparisons were performed separately within each mESC clone. Dots indicate the mean TE difference, and error bars represent 95% confidence intervals. Comparisons are shown for (A) genes with ASE versus genes without ASE (mean TE difference: −0.563 to −0.275; *p* < 0.008), (B) genes with consistent ASE versus genes without ASE (−0.781 to −0.489; *p* < 0.0003), (C) genes with variable ASE versus genes without ASE (−0.368 to −0.041; *p* < 0.002), and (D) genes classified as non-ASE overall but exhibiting clone-specific allelic imbalance versus genes without allelic imbalance in the corresponding clone (−0.283 to −0.080; *p* < 0.0005).

**S16 Figure.**
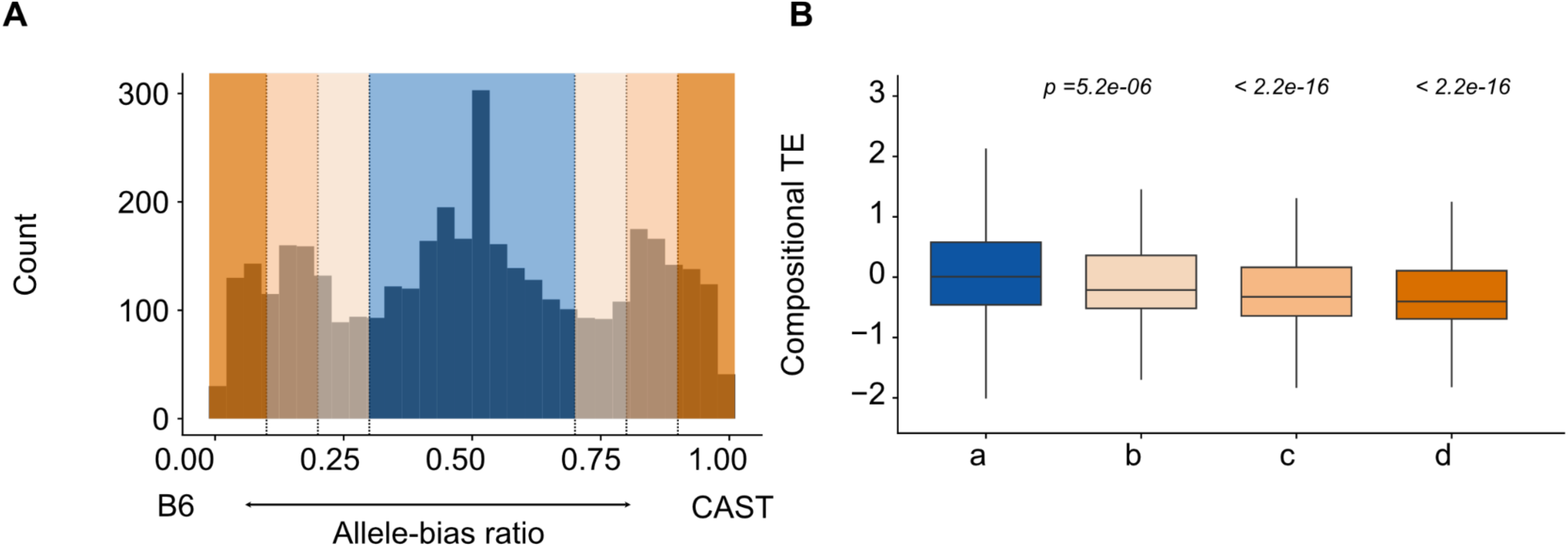
Compositional TE of mESC genes across increasing allelic-bias ratios. (A) Distribution of allelic-bias ratios for genes with ASE and expression-matched genes without ASE. Genes were classified based on their allelic bias: Group (a), 0.3 < allele bias ratio < 0.7; Group (b), allelic bias ratio < 0.3 or > 0.7; Group (c), allelic bias ratio < 0.2 or > 0.8; Group (d), allelic bias ratio < 0.1 or > 0.9. (B) Compositional TE was compared after matching RNA expression levels between each ASE group and the non-ASE group. P-values were calculated using the Wilcoxon rank-sum test.

**S17 Figure.**
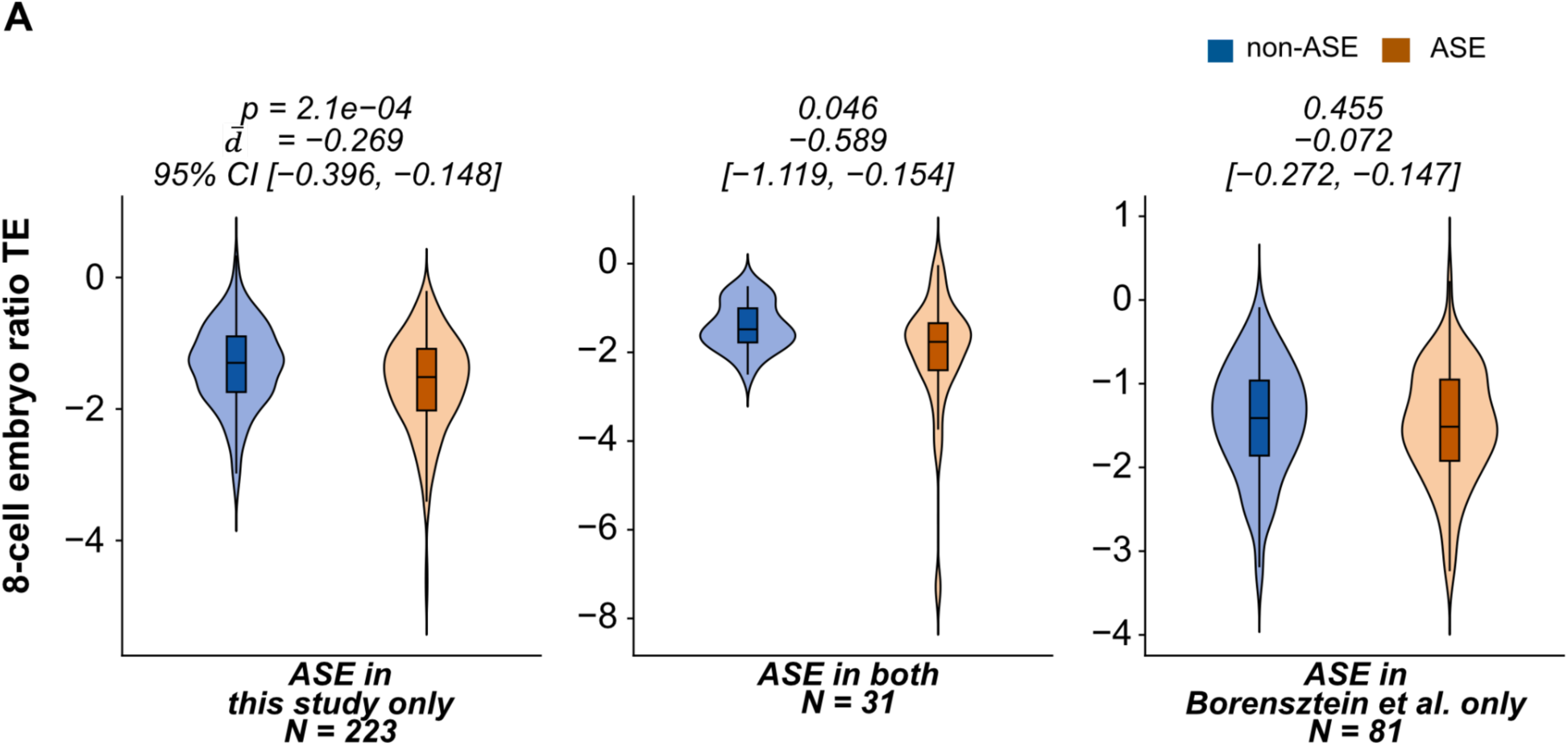
Ratio TE comparison between ASE defined from independent embryo datasets. Ratio TE of genes exhibiting ASE defined from two independent 8-cell embryo RNA-seq datasets was evaluated using our embryo TE measurements. Because the published Borensztein et al. (2017) dataset did not include ribosome profiling, TE was calculated using our RNA-seq and ribosome profiling dataset. Genes were grouped into three categories: ASE in our dataset only, Borensztein only, genes with ASE in both datasets. P-values were calculated using the Wilcoxon rank-sum test by comparing each group.

**S18 Figure.**
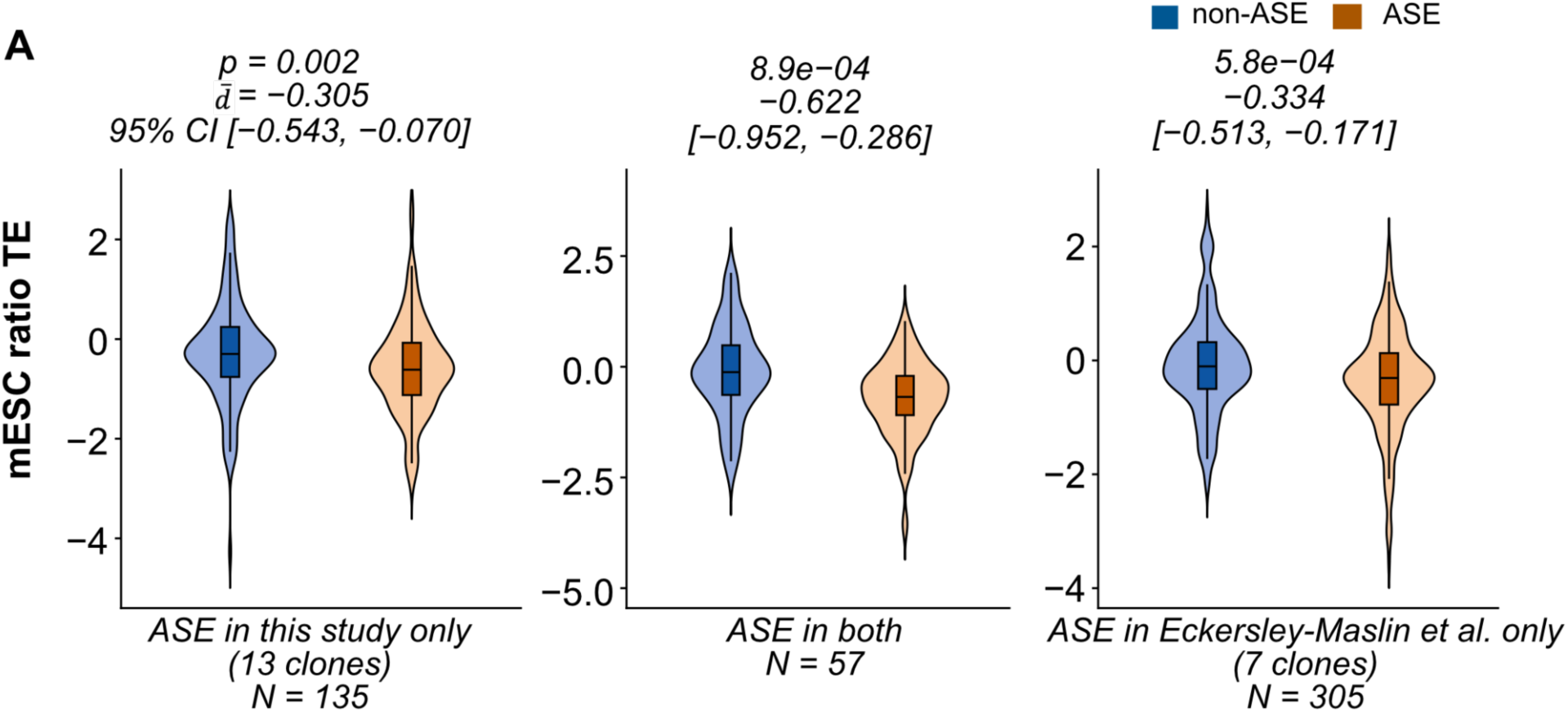
TE comparison between ASE defined from independent mESC datasets. (A) Ratio TE comparison of genes exhibiting ASE defined from two independent mESC clone RNA-seq datasets. Genes with ASE were identified from our mESC clone dataset and the published dataset by Eckersley-Maslin et al. (2014). Genes were grouped into three categories: ASE identified only in our dataset, ASE identified only in the Eckersley-Maslin dataset, overlapping genes with ASE identified in both datasets. For each ASE group, genes without ASE were selected by matching RNA expression levels to the corresponding genes exhibiting ASE. Ratio TE was calculated using our matched RNA-seq and ribosome profiling data. P-values were calculated using the Wilcoxon rank-sum test comparing each ASE group with its RNA expression-matched non-ASE group.

**S19 Figure.**
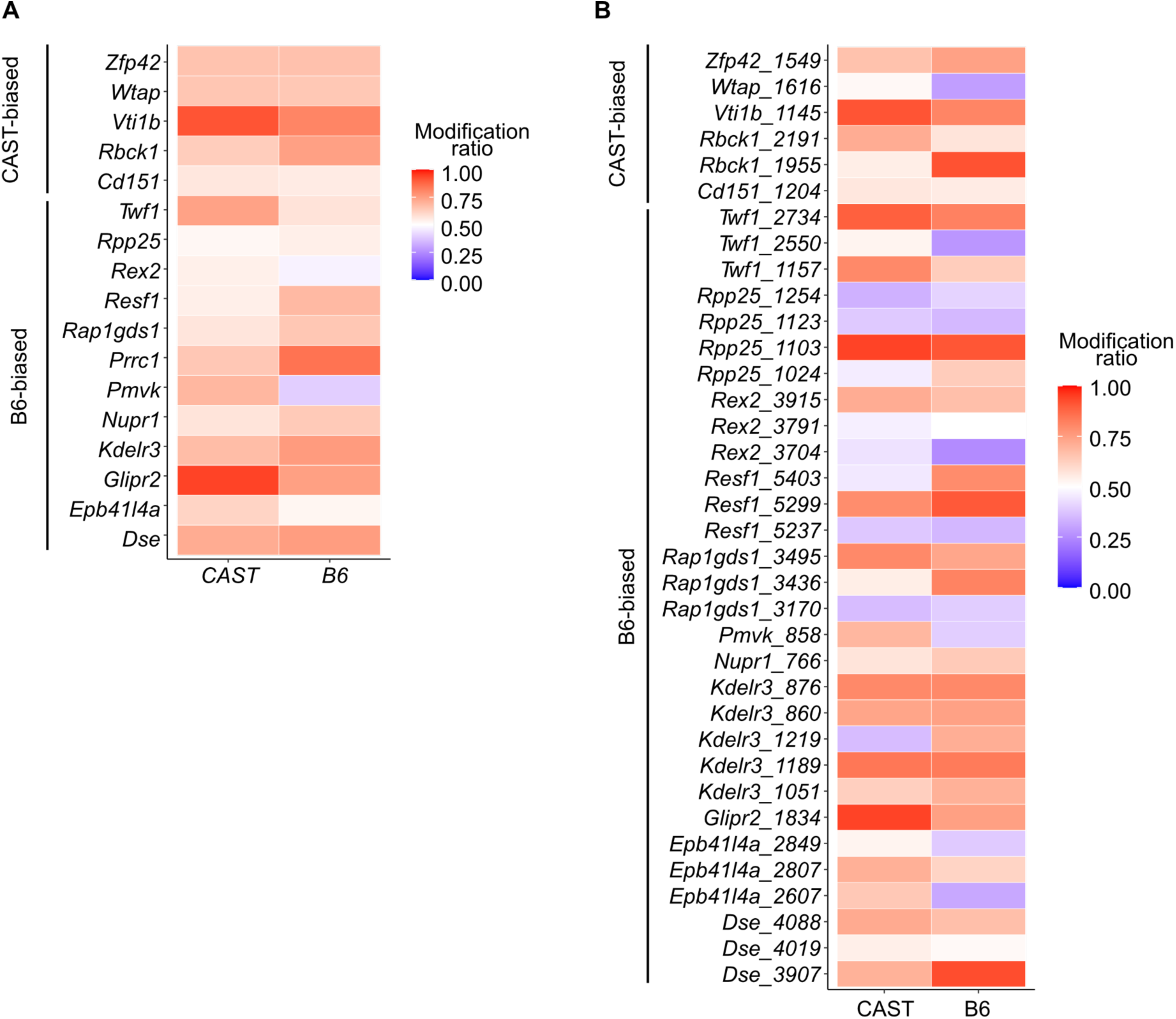
Allele-specific m6A modification levels in genes with and without ASE. Allele-specific m6A modification levels were quantified by measuring the m6A modification ratio separately for the B6 and CAST alleles of genes classified as ASE or non-ASE. (A) Sum of all allele-specific m6A modification ratios across each transcript. (B) Sum of allele-specific m6A modification ratios within the 3′ UTR. No significant differences in allele-specific m6A modification levels were observed between genes with and without ASE.

**S20 Table.**
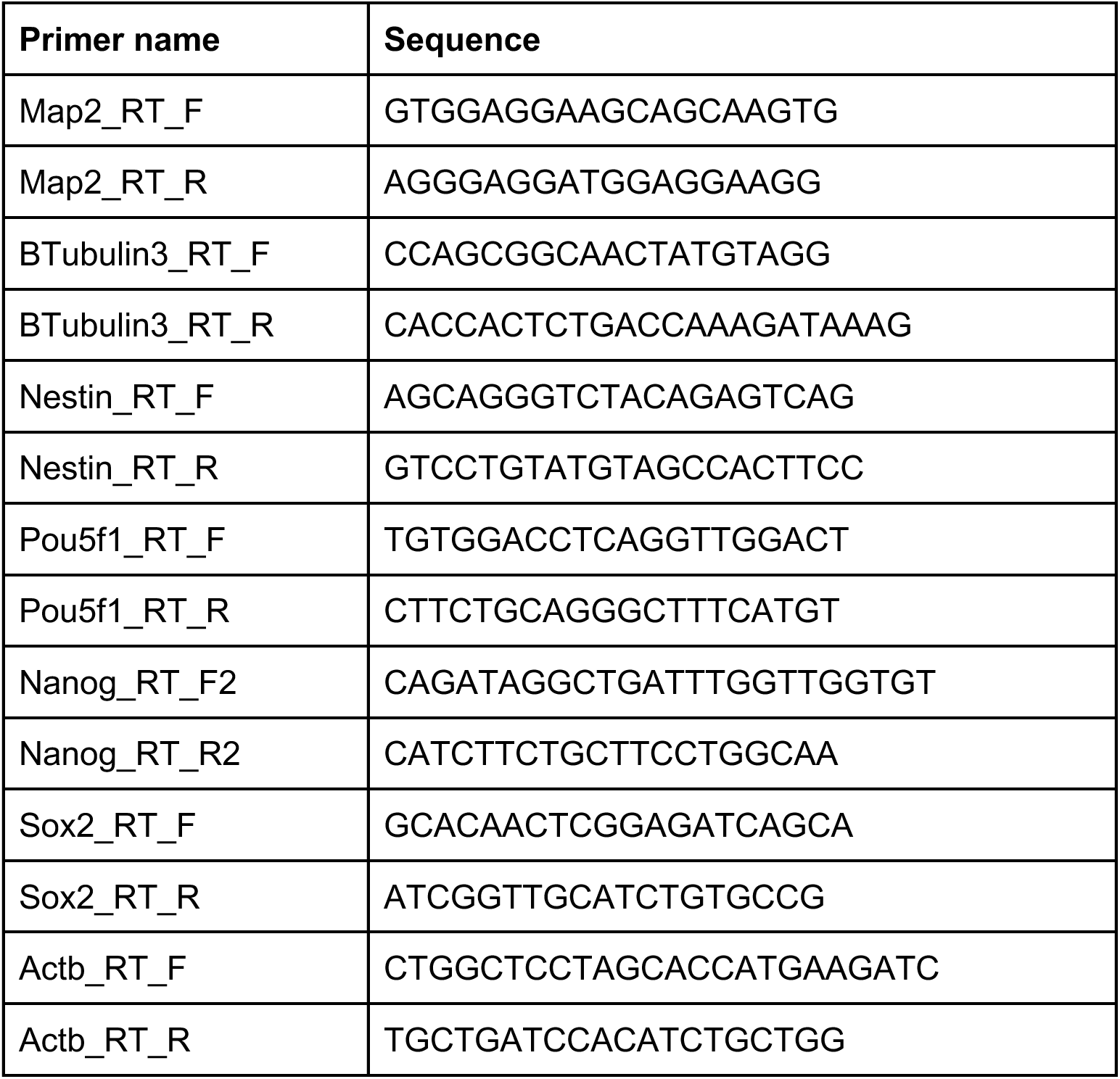
Primer sequences for qRT-PCR.

**S21 Table.** Total mapped read counts. Mapped read counts from RNA-seq and ribosome profiling libraries. For RNA-seq, reads were mapped to the transcriptome. For ribosome profiling, reads between 15 and 50 nucleotides in length were retained and mapped to the transcriptome.

| Sample name | Transcriptome mapped read count |
| --- | --- |
| Ribo_mESC_clone_A | 644,528 |
| Ribo_mESC_clone_B | 955,992 |
| Ribo_mESC_clone_C | 533,151 |
| Ribo_mESC_clone_D | 2,126,758 |
| Ribo_mESC_clone_E | 888,567 |
| Ribo_mESC_clone_F | 1,745,667 |
| Ribo_mESC_clone_G | 379,044 |
| Ribo_mESC_clone_H | 661,145 |
| Ribo_mESC_clone_I | 1,779,776 |
| Ribo_mESC_clone_J | 228,200 |
| Ribo_mESC_clone_K | 3,448,470 |
| Ribo_mESC_clone_L | 2,917,283 |
| Ribo_mESC_clone_M | 5,195,797 |
| RNA_mESC_clone_A | 5,183,551 |
| RNA_mESC_clone_B | 4,951,735 |
| RNA_mESC_clone_C | 2,114,068 |
| RNA_mESC_clone_D | 3,733,372 |
| RNA_mESC_clone_E | 3,152,604 |
| RNA_mESC_clone_F | 4,785,600 |
| RNA_mESC_clone_G | 4,578,365 |
| RNA_mESC_clone_H | 4,201,009 |
| RNA_mESC_clone_I | 3,659,820 |
| RNA_mESC_clone_J | 5,750,488 |
| RNA_mESC_clone_K | 3,752,508 |
| RNA_mESC_clone_L | 4,083,219 |
| RNA_mESC_clone_M | 4,743,627 |
| Ribo_NPC_clone_A | 185,317 |
| Ribo_NPC_clone_B | 213,684 |
| Ribo_NPC_clone_C | 200,514 |
| Ribo_NPC_clone_D | 278,218 |
| Ribo_NPC_clone_E | 44,108 |
| RNA_NPC_clone_A | 6,275,571 |
| RNA_NPC_clone_B | 4,968,805 |
| RNA_NPC_clone_C | 5,466,725 |
| RNA_NPC_clone_D | 6,349,630 |
| RNA_NPC_clone_E | 6,476,725 |
| Ribo_8cell_BxC_A | 103,233 |
| Ribo_8cell_BxC_B | 181,355 |
| Ribo_8cell_BxC_C | 135,312 |
| Ribo_8cell_BxC_D | 111,441 |
| Ribo_8cell_CxB_A | 330,093 |
| Ribo_8cell_CxB_B | 267,022 |
| Ribo_8cell_CxB_C | 188,690 |
| Ribo_8cell_CxB_D | 198,227 |
| RNA_8cell_BxC_A_Smart-seq3 | 5,499,357 |
| RNA_8cell_BxC_B_Smart-seq3 | 5,706,972 |
| RNA_8cell_BxC_C_Smart-seq3 | 6,498,875 |
| RNA_8cell_BxC_D_Smart-seq3 | 4,303,088 |
| RNA_8cell_CxB_A_Smart-seq3 | 4,431,276 |
| RNA_8cell_CxB_B_Smart-seq3 | 7,338,891 |
| RNA_8cell_CxB_C_Smart-seq3 | 8,062,311 |
| RNA_8cell_CxB_A_NEB | 30,712,799 |
| RNA_8cell_CxB_B_NEB | 21,430,381 |
| RNA_8cell_CxB_C_NEB | 2,172,205 |
| RNA_8cell_CxB_D_NEB | 25,028,367 |
| RNA_adult_kidney_rep1 | 5,181,477 |
| RNA_adult_kidney_rep2 | 5,437,553 |
| RNA_adult_kidney_rep3 | 5,823,463 |
| Ribo_adult_kidney_rep1 | 2,432,901 |
| Ribo_adult_kidney_rep2 | 3,324,403 |
| Ribo_adult_kidney_rep3 | 2,693,617 |
| RNA_adult_liver_rep1 | 11,025,320 |
| RNA_adult_liver_rep2 | 14,672,440 |
| RNA_adult_liver_rep3 | 19,249,570 |
| Ribo_adult_liver_rep1 | 1,288,078 |
| Ribo_adult_liver_rep2 | 1,728,411 |
| Ribo_adult_liver_rep3 | 1,765,718 |
| RNA_adult_lung_rep1 | 13,705,615 |
| RNA_adult_lung_rep2 | 6,523,370 |
| RNA_adult_lung_rep3 | 5,181,477 |
| Ribo_adult_lung_rep1 | 4,248,536 |
| Ribo_adult_lung_rep2 | 4,323,541 |
| Ribo_adult_lung_rep3 | 3,879,729 |

**S22 Figure.**
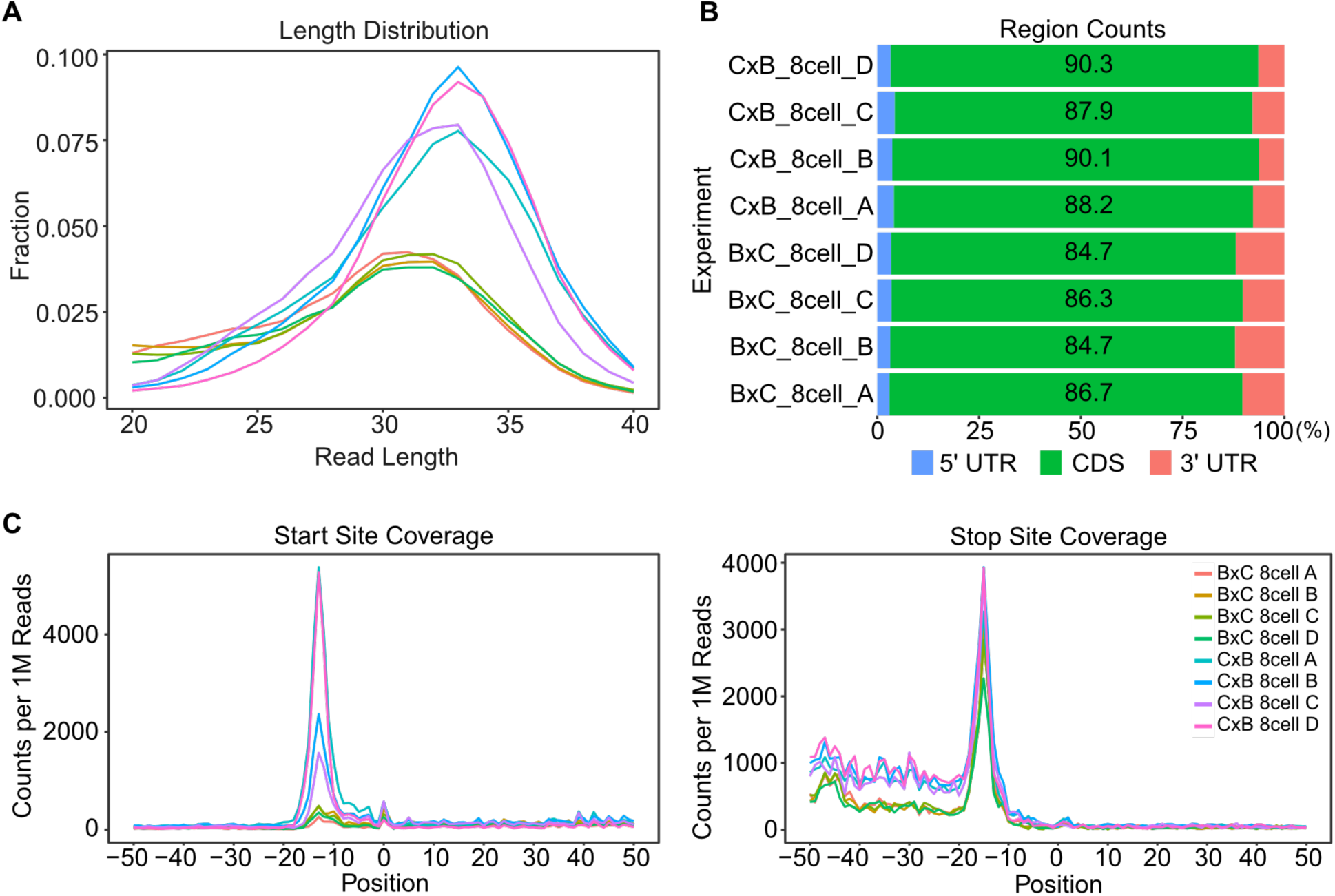
Quality control of 8-cell reciprocal cross embryo ribosome profiling data. (A) Ribosome profiling read length distribution. (B) Distribution of reads across genomic regions. (C–D) Read density at annotated start codons (C) and stop codons (D).

